# TGFβ Suppresses Type 2 Immunity to Cancer

**DOI:** 10.1101/2020.03.04.977629

**Authors:** Ming Liu, Fengshen Kuo, Kristelle J. Capistrano, Davina Kang, Briana G. Nixon, Wei Shi, Chun Chou, Mytrang H. Do, Shun Li, Efstathios G. Stamatiades, Yingbei Chen, James J. Hsieh, A. Ari Hakimi, Ichiro Taniuchi, Timothy A. Chan, Ming O. Li

## Abstract

The immune system employs two distinct defense strategies against infections: pathogen elimination typified by type 1 immunity, and pathogen containment exemplified by type 2 immunity in association with tissue repair. Akin to infectious diseases, cancer progresses with cancer cell acquisition of microorganism-like behavior propagating at the expense of the host. While immunological mechanisms of cancer cell elimination are well defined, whether immune-mediated cancer cell containment can be induced is poorly understood. Here we show that ablation of transforming growth factor-β receptor II (TGFβRII) in CD4^+^ T cells promotes tumor tissue healing and halts cancer progression. Notably, the restorative response is dependent on the T helper 2 cytokine IL-4 fortifying vasculature organization that spares only proximal layers of cancer cells from hypoxia, nutrient starvation and death. Thus, type 2 immunity represents an effective cancer defense mechanism, and TGFβ signaling in helper T cells may be targeted for novel cancer immunotherapy.

## Text

In response to invading pathogens, vertebrate immune system employs two discrete strategies of host defense: pathogen elimination typified by type 1 immunity following innate immune recognition of prokaryotic microorganisms^1^, and pathogen containment associated with tissue repair manifested by type 2 immunity following the invasion of metazoan parasites that inflict tissue damage^2^. Similar to infectious diseases, cancer imposes insults to the host with cancer cell acquisition of microorganism-like behavior of clonal expansion. Indeed, cancer cell-targeted type 1 immunity mediated by CD8^+^ cytotoxic T lymphocytes (CTLs), cytotoxic innate lymphocytes and innate-like T cells represent host protective mechanisms of cancer cell elimination^3-5^. Notwithstanding, the host impact of cancer cells is much more intimate than that of microbes, and cancer pathogenicity, in particular that of solid tumors, is dependent on cancer cell coercing a supportive environment that promotes tumor encroachments leading to the impairment of tissue function^6-9^. Notably, tumor tissues resemble active wounds, and are considered as “wounds that do not heal”^10^, implying that cancer microenvironment-targeted tissue repair responses could restrain tumor progression.

Dissimilar from infectious diseases, cancer arises from malignant transformation of self-cells. Consequently, immunological self-tolerance can be maladapted to repress cancer-elicited immune responses. Indeed, multiple T cell self-tolerance regulators including Foxp3^+^ regulatory T (Treg) cells, co-inhibitory receptors PD-1 and CTLA-4, and the regulatory cytokine β) suppress immune control of tumor development. Depletion of tumor-infiltrating Treg cells via genetic approaches or administration of anti-PD-1 and anti-CTLA-4 result in CTL-dependent cancer cell elimination in animal models and patients^16,17^. In transgenic mouse models of cancer, blockade of TGFβ signaling in T cells suppresses tumor development^18^, which is correlated with expansion of tumor-reactive CTLs that exhibit enhanced cancer cell killing *in vitro*. Nonetheless, as TGFβ exerts pleiotropic effects on multiple lineages of T cells^19^, its functional targets *in vivo* and the underlying mechanisms of cancer regulation remain obscure.

### TGFβ acts on CD4 T cells to foster tumor growth

As CTLs are established effectors of cancer cell elimination, we set forth by examining whether TGF β directly suppressed CTL-mediated cancer surveillance in the MMTV-PyMT (PyMT) model of breast cancer. Mice carrying a floxed allele of the *Tgfbr2* gene (*Tgfbr2^fl/fl^*) encoding the TGF β receptor II (TGFβRII) were crossed with CD8^Cre^ transgenic mice, which were further bred to the PyMT background. Loss of TGFβRII was observed specifically in CD8^+^ T cells from CD8^Cre^*Tgfbr2^fl/fl^*PyMT mice (Fig. 1a), which led to enhanced effector/memory differentiation of CD8^+^ T cell in the tumor-draining lymph nodes (Extended Data Fig. 1). Increased expression of the cytolytic enzyme Granzyme B among PD-1-expressing CD8^+^ T cells was also detected in the tumor (Fig. 1b). Surprisingly, mammary tumor growth was not suppressed in CD8^Cre^*Tgfbr2^fl/fl^*PyMT mice (Fig. 1c). These observations imply that additional immunosuppressive mechanisms such as those mediated by PD-1 whose expression was unaffected in tumor-infiltrating CD8^+^ T cells (Fig. 1b), may compensate for TGFβ signaling blockade for CTL repression as suggested by recent studies^20-23^.

**Figure 1.**
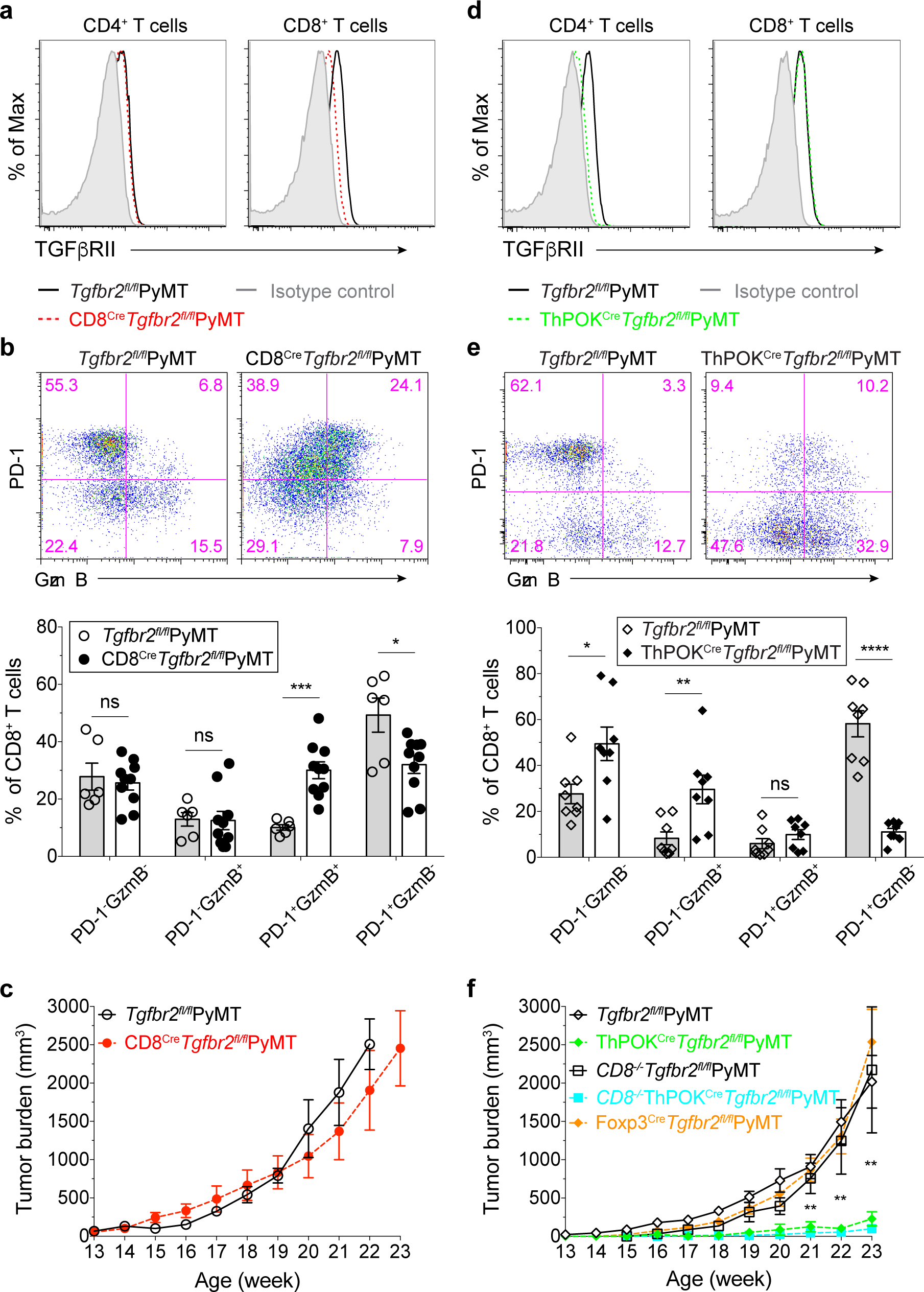
Blockade of TGFβ signaling in CD4^+^ T cells suppresses tumor development independent of CD8^+^ T cells **a,** Transforming growth factor-β receptor II (TGFβRII) expression on CD4^+^ T cells and CD8^+^ T cells from the tumor-draining lymph nodes of *Tgfbr2^fl/fl^*PyMT and CD8^Cre^*Tgfbr2^fl/fl^*PyMT mice. **b,** Representative flow cytometry plots and statistical analyses of programmed cell death protein 1 (PD-1) and granzyme B (GzmB) expression in tumor-infiltrating CD8^+^ T cells from *Tgfbr2^fl/fl^*PyMT and CD8^Cre^*Tgfbr2^fl/fl^*PyMT mice. **c,** Tumor measurements of *Tgfbr2^fl/fl^*PyMT (n=8) and CD8^Cre^*Tgfbr2^fl/fl^*PyMT (n=7) mice. **d**, TGFβRII expression on CD4^+^ T cells and CD8^+^ T cells from the tumor-draining lymph nodes of *Tgfbr2^fl/fl^*PyMT and ThPOK^Cre^*Tgfbr2^fl/fl^*PyMT mice. **e,** Representative flow cytometry plots and statistical analyses of PD-1 and GzmB expression in tumor-infiltrating CD8^+^ T cells from *Tgfbr2^fl/fl^*PyMT and ThPOK^Cre^*Tgfbr2^fl/fl^*PyMT mice. **f,** Tumor measurements of *Tgfbr2^fl/fl^*PyMT (n=7), ThPOK^Cre^*Tgfbr2^fl/fl^*PyMT (n=5), *CD8^-/-^ Tgfbr2^fl/fl^*PyMT (n=3), *CD8^-/-^*ThPOK^Cre^*Tgfbr2^fl/fl^*PyMT (n=5) and Foxp3^Cre^*Tgfbr2^fl/fl^*PyMT (n = 3) mice. All data are shown as mean ± SEM. *: P<0.05; **: P<0.01; ***: P<0.001; ****: P<0.0001; and ns: not significant.

To investigate whether TGFβ acted on CD4^+^ T cells to indirectly suppress CTL-dependent cancer surveillance, we crossed *Tgfbr2^fl/fl^*PyMT mice with ThPOK^Cre^ mice that predominantly targeted CD4^+^, but not CD8^+^, innate-like or TCRγδ T cells (Fig. 1d and Extended Data Fig. 2a). Compared to control *Tgfbr2^fl/fl^*PyMT mice, ThPOK^Cre^*Tgfbr2^fl/fl^*PyMT mice exhibited enhanced effector/memory differentiation of conventional CD4^+^ T cells and Treg cells, but not CD8^+^ T cells, in the tumor-draining lymph nodes (Extended Data Fig. 2b). Yet, tumor-infiltrating CD8^+^ T cells expressed higher levels of Granzyme B and lower levels of PD-1 (Fig. 1e). Such a phenotypic change was observed in T cell-specific TGFβ1-deficeint mice that resist tumor growth^18^. Indeed, profound inhibition of tumor progression was observed in ThPOK^Cre^*Tgfbr2^fl/fl^*PyMT mice (Fig. 1f). To determine whether the altered CTL response accounted for the tumor growth phenotype, we crossed ThPOK^Cre^*Tgfbr2^fl/fl^*PyMT mice to the CD8-deficient background. Unexpectedly, tumor repression was unchanged in the absence of CD8^+^ T cells (Fig. 1f), while depletion of CD4^+^ T cells restored tumor growth (Extended Data Fig. 2c-2d). These findings suggest that CD4^+^ T cells are a prominent target of TGFβ in cancer immune tolerance control, and the reduced tumor growth triggered by TGFβ cells is not mediated by CTLs.

**Figure 2.**
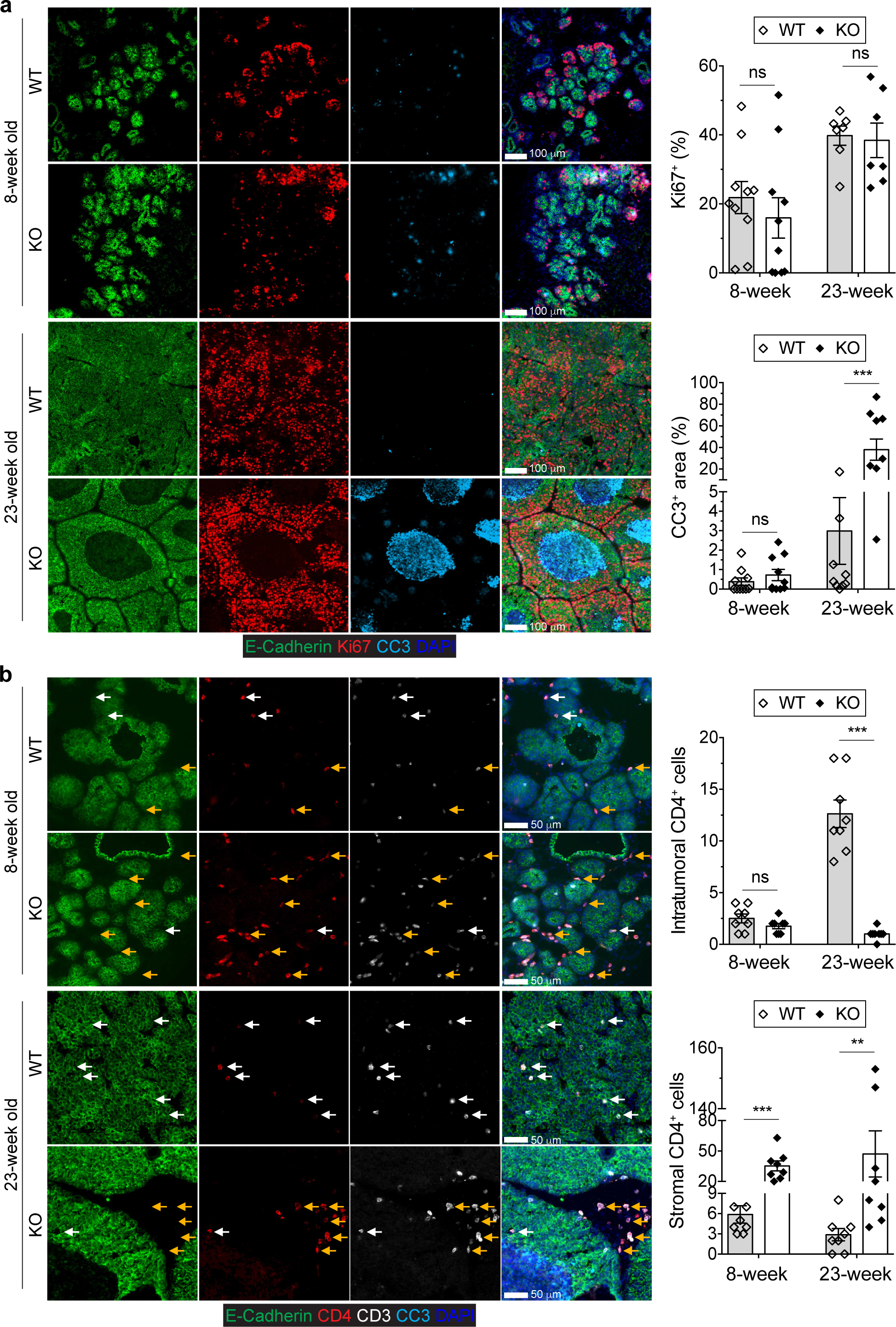
Blockade of TGFβ signaling in CD4^+^ T cells triggers cancer cell death and stroma localization of CD4^+^ T cells **a,** Representative immunofluorescence images and statistical analyses of E-Cadherin (green), Ki67 (red) and cleaved Caspase 3 (CC3, cyan) expression in mammary tumor tissues from 8- and 23-week-old *Tgfbr2^fl/fl^*PyMT (wild-type, WT) and ThPOK^Cre^*Tgfbr2^fl/fl^*PyMT (knockout, KO) mice. The percentage of Ki67^+^E-Cadherin^+^ cells over total E-Cadherin^+^ epithelial cells was calculated from multiple 0.02 mm^2^ regions (n=10 and 7 for WT and KO tumor tissues, respectively). The percentage of CC3^+^ areas over total E-Cadherin^+^ areas was calculated from multiple 0.02 mm^2^ regions (n=10 for WT and KO tumor tissues). **b,** Representative immunofluorescence images and statistical analyses of E-Cadherin (green), CD4 (red), CD3 (white) and CC3 (cyan) expression in mammary tumor tissues from 8- and 23-week-old WT and KO mice. Intratumoral (white arrows) and stromal (yellow arrows) CD4^+^ T cells were counted from multiple 0.1 mm^2^ regions (n=8 for WT and KO tumor tissues). All data are shown as mean ± SEM. **: P<0.01; ***: P<0.001; and ns: not significant.

### Cancer cell death occurs with immune exclusion

To define the underlying mechanisms of cancer suppression in ThPOK^Cre^*Tgfbr2^fl/fl^*PyMT mice, we assessed proliferation and death of cancer cells by the expression of Ki67 and cleaved Caspase 3 (CC3), respectively. Ki67 was expressed in about 20% and 40% mammary epithelial cells in tumors from 8-week-old and 23-week-old *Tgfbr2^fl/fl^*PyMT mice, which was unaffected in ThPOK^Cre^*Tgfbr2^fl/fl^*PyMT mice (Fig. 2a). In contrast, blockade of TGFβ signaling in CD4^+^ T cells resulted in an approximate 13-fold increase of CC3-positive cells at 23-week of age (Fig. 2a). Notably, dying cancer cells had a clustered distribution pattern (Fig. 2a), which was not observed in CD8^Cre^*Tgfbr2^fl/fl^*PyMT mice (Extended Data Fig. 3a), but was preserved in ThPOK^Cre^*Tgfbr2^fl/fl^*PyMT mice crossed onto the CD8-deficient background (Extended Data Fig. 3b). These observations imply that TGFβRII-deficient CD4^+^ T cells inhibit tumor progression through the induction of cancer cell death.

**Figure 3.**
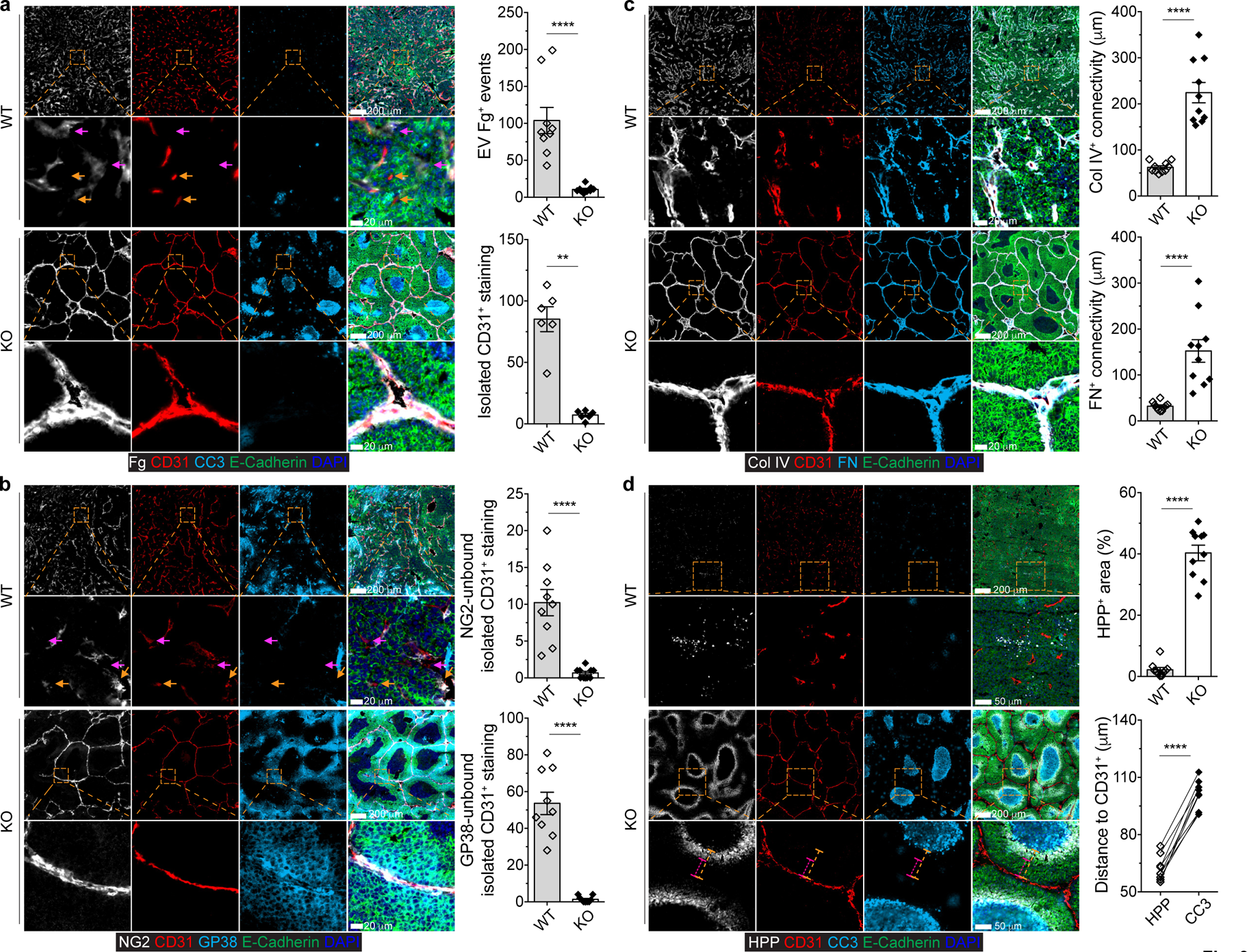
Blockade of TGFβ signaling in CD4^+^ T cells promotes tumor tissue healing, vessel organization, hypoxia and cancer cell death **a,** Representative immunofluorescence images of fibrinogen (Fg, white), CD31 (red), cleaved Caspase 3 (CC3, cyan) and E-Cadherin (green) in comparable individual tumors with sizes around 8×8 mm in length and width from 23-week-old *Tgfbr2^fl/fl^*PyMT (wild-type, WT) and ThPOK^Cre^*Tgfbr2^fl/fl^*PyMT (knockout, KO) mice. Extravascular (EV) Fg deposition events (magenta arrows) were calculated from multiple 1 mm^2^ regions (n=9 for WT and KO tumor tissues). Isolated CD31^+^ staining (yellow arrows) was counted from multiple 1 mm^2^ regions (n=6 for WT and KO tumor tissues). **b,** Representative immunofluorescence images of NG2 (white), CD31 (red), GP38 (cyan) and E-Cadherin (green) in comparable individual tumors with sizes around 8×8 mm in length and width from 23-week-old WT and KO mice. NG2-unbound (magenta arrows) or GP38-unbound (yellow arrows) isolated CD31^+^ staining was counted from multiple 1 mm^2^ regions (n=9 for WT and KO tumor tissues). **c,** Representative immunofluorescence images of collagen IV (Col IV, white), CD31 (red), fibronectin (FN, cyan) and E-Cadherin (green) in comparable individual tumors with sizes around 8×8 mm in length and width from WT and KO mice. The average continuous lengths of Col IV and FN were measured in multiple 1 mm^2^ regions (n=9 for WT and KO tumor tissues). **d,** Representative immunofluorescence images of Hypoxic probe (HPP, white), CD31 (red), CC3 (cyan) and E-Cadherin (green) in comparable individual tumors with sizes around 8×8 mm in length and width from WT and KO mice. The percentage of HPP^+^E-Cadherin^+^ regions over E-Cadherin^+^ epithelial regions was calculated from multiple 1 mm^2^ regions (n=9 for WT and KO tumor tissues). The shortest distance of HPP^+^ regions (magenta dashed lines) or CC3^+^ regions (yellow dashed lines) to CD31^+^ endothelial cells was measured in tumor tissues from KO mice (n=9). The dashed boxes coupled with dashed lines show high magnification of selected tissue regions. All data are shown as mean ± SEM. **: P<0.01; ****: P<0.0001.

The lack of a role for CTLs in tumor repression in ThPOK^Cre^*Tgfbr2^fl/fl^*PyMT mice prompted us to investigate whether CD4^+^ T cells might directly eradicate cancer cells as suggested by recent studies^24,25^. To this end, we examined the localization of CD4^+^ T cells by immunofluorescence staining. Tumor progression was associated with an approximate 5-fold increase of intratumoral CD4^+^ T cells between 8-week-old and 23-week-old *Tgfbr2^fl/fl^*PyMT mice, while stromal CD4^+^ T cells were reduced (Fig. 2b). In contrast, blockade of TGFβ signaling in CD4^+^ T cells led to approximate 6- and 16-fold increase of stromal CD4^+^ T cells in 8-week-old and 23-week-old mice, respectively, while intratumoral CD4^+^ T cells were substantially reduced in 23-week-old ThPOK^Cre^*Tgfbr2^fl/fl^*PyMT mice (Fig. 2b). Furthermore, few intratumoral CD4^+^ T cells were localized distant from the cancer cell death region (Fig. 2b), suggesting against direct cancer cell killing by TGFβRII-deficient CD4^+^ T cells. To examine whether the preferential tumor stromal localization of CD4^+^ T cells in ThPOK^Cre^*Tgfbr2^fl/fl^*PyMT mice was applicable to other hematopoietic lineage cells, we performed immunofluorescence staining with the pan-leukocyte marker CD45. In contrast to the dominant tumor parenchyma localization of CD45^+^ cells in *Tgfbr2^fl/fl^*PyMT mice, leukocytes were mostly localized in the tumor stroma of ThPOK^Cre^*Tgfbr2^fl/fl^*PyMT mice (Extended Data Fig. 4). The immune cell exclusion phenotype suggests that TGFβRII-deficient CD4^+^ T cells unlikely induce cancer cell death directly or indirectly via another leukocyte effector.

**Figure 4.**
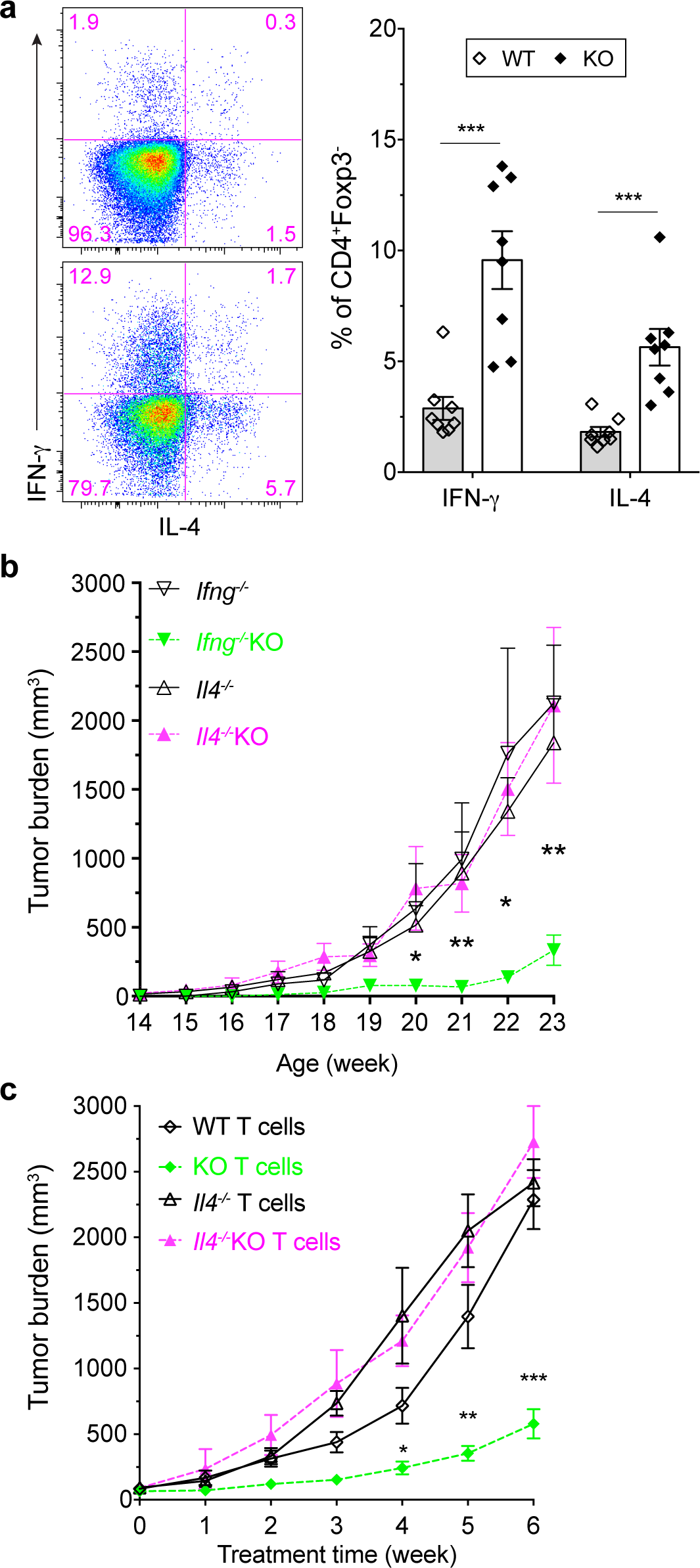
Anti-tumor immunity triggered by TGFβ signaling blockade in CD4^+^ T cells is dependent on IL-4, but not IFN-**γ a,** Representative flow cytometry plots and statistical analyses of IL-4 and IFN-γ expression in CD4^+^Foxp3^-^ T cells from the tumor-draining lymph nodes of 23-week-old *Tgfbr2^fl/fl^*PyMT (wild-type, WT) and ThPOK^Cre^*Tgfbr2^fl/fl^*PyMT (knockout, KO) mice. **b**, Tumor measurements of *Ifng*^-/-^ *Tgfbr2^fl/fl^*PyMT (*Ifng*^-/-^, n=3), *Ifng*^-/-^ThPOK^Cre^*Tgfbr2^fl/fl^*PyMT (*Ifng*^-/-^KO, n=6), *Il4^-/-^Tgfbr2^fl/fl^*PyMT (*Il4^-/-^*, n=4) and *Il4*^-/-^ThPOK^Cre^*Tgfbr2^fl/fl^*PyMT (*Il4^-/-^*KO, n=4) mice. **c,** 16 to 17-week-old PyMT mice bearing 5×5 mm tumors were transferred with CD4^+^CD25^-^ T cells from *Tgfbr2^fl/fl^* (WT, n=4), ThPOK^Cre^*Tgfbr2^fl/fl^* (KO, n=4), *Il4^-/-^Tgfbr2^fl/fl^* (*Il4^-/-^, n=3*) and *Il4^-/-^*ThPOK^Cre^*Tgfbr2^fl/fl^* (*Il4^-/-^*KO, n=3) mice on a weekly basis for 6 weeks. Tumor burden was measured and plotted. All data are shown as mean ± SEM. *: P<0.05; **: P<0.01; ***: P<0.001.

### Vessel organization triggers cancer cell death

The preferential stromal localization of TGFβRII-deficient CD4^+^ T cells suggests that they may regulate the host to endure the negative impact of a growing tumor with cancer cell death being a secondary outcome. In line with previous studies^10^, the fast growing tumor in *Tgfbr2^fl/fl^*PyMT mice exhibited extensive extravascular deposition of fibrinogen (Fig. 3a), indicative of vasculature damage. In contrast, fibrinogen was predominantly intravascular in ThPOK^Cre^*Tgfbr2^fl/fl^*PyMT mice (Fig. 3a). Perfusion with a biotinylation probe confirmed an overt leaky vasculature in tumors from *Tgfbr2^fl/fl^*PyMT, but not ThPOK^Cre^*Tgfbr2^fl/fl^*PyMT mice (Extended Data Fig. 5a). The vasculature damage phenotype in *Tgfbr2^fl/fl^*PyMT mice co-occurred with an irregularly shaped and bluntly ended microvasculature manifested by the isolated staining of the endothelium marker CD31 (Fig. 3a). Strikingly, although the vessel density was unaffected (Extended Data Fig. 5b), tumor vasculature was much more organized in ThPOK^Cre^*Tgfbr2^fl/fl^*PyMT mice with less isolated CD31^+^ endothelial cell staining (Fig. 3a).

**Figure 5.**
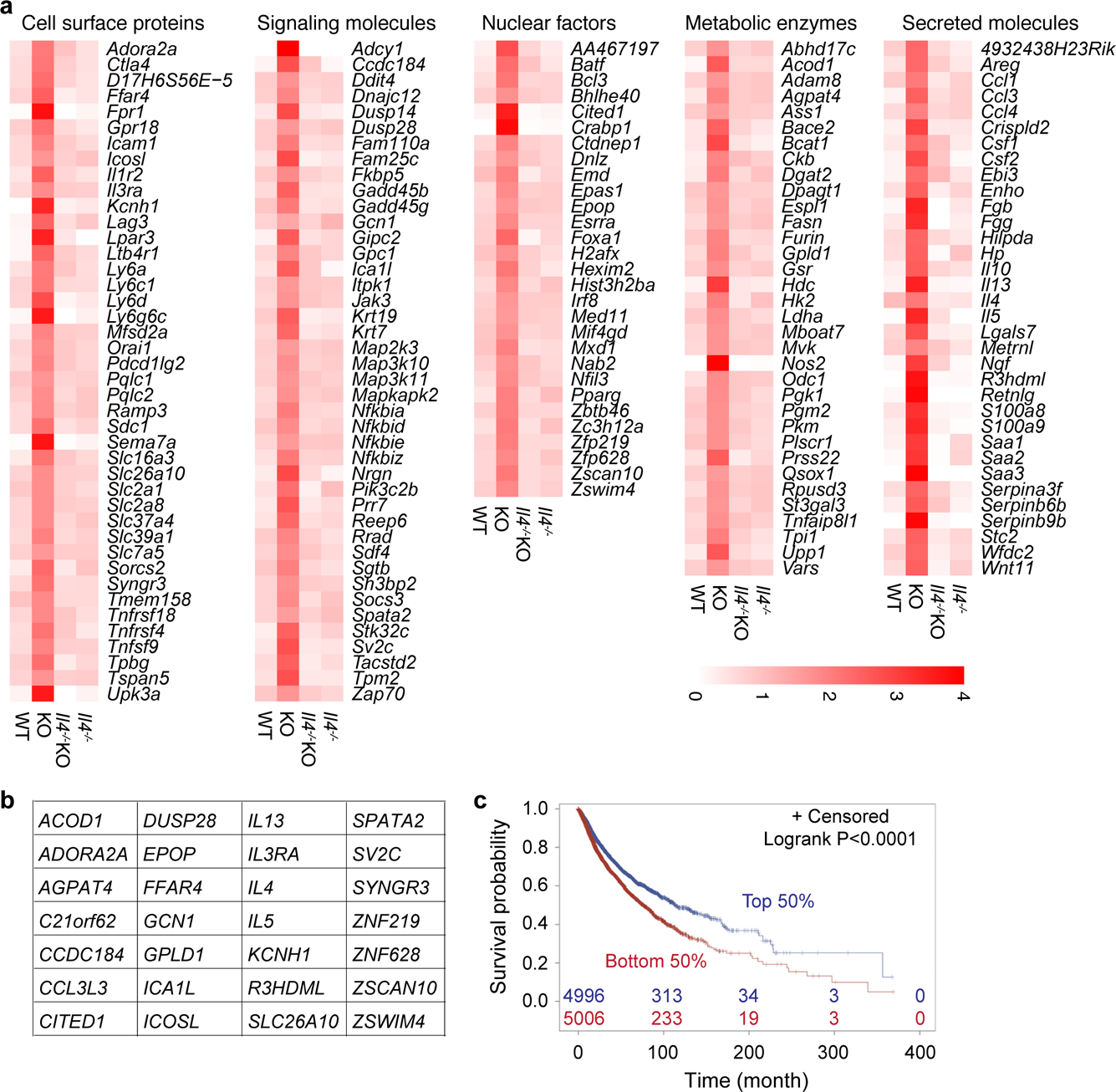
A TGFβ-regulated IL-4-dependent gene expression signature stratifies cancer patients for survival probability **a,** The transcriptome of tumor-infiltrating CD4^+^CD25^-^ T cells from 23-week-old *Tgfbr2^fl/fl^*PyMT (wild-type, WT), ThPOK^Cre^*Tgfbr2^fl/fl^*PyMT (knockout, KO), *Il4^-/-^Tgfbr2^fl/fl^*PyMT (*Il4^-/-^*) and *Il4^-/-^* ThPOK^Cre^*Tgfbr2^fl/fl^*PyMT (*Il4^-/-^*KO) mice were assessed by RNA sequencing. The mean expression FPKM value of differentially expressed genes (DEGs) that were upregulated in KO compared to WT or *Il4^-/-^*KO samples was calculated and plotted. The ratio between mean FPKM value of one group and average of mean FPKM values of all 4 groups was further calculated. Representative DEGs were grouped based on the localization and function of their encoded proteins. **b,** A Th2 gene expression signature was generated through comparing KO vs. WT and KO vs. *Il4^-/-^*KO in the upregulated direction after additional filtering. **c,** The Th2 gene expression signature was used to perform survival analysis in TCGA Pan cancer cohort data and survival curves were plotted for the signature high group (top 50%) and low group (bottom 50%). The corresponding censored patient numbers are included in major time points.

These observations demonstrate that while tumors from *Tgfbr2^fl/fl^*PyMT mice inflict chronic tissue damage resembling “wounds that do not heal”^10^, tumors from ThPOK^Cre^*Tgfbr2^fl/fl^*PyMT mice are maintained in a healed state associated with an organized vasculature.

Pericytes are mesenchyme-derived cells that enwrap and stabilize capillaries and control perfusion^26^. Notably, while approximate 12% endothelial cells with isolated CD31^+^ staining in tumors from *Tgfbr2^fl/fl^*PyMT mice were not bound by NG2^+^ pericytes (Fig. 3a-3b), the endothelium was tightly ensheathed by pericytes in tumors from ThPOK^Cre^*Tgfbr2^fl/fl^*PyMT mice (Fig. 3b). Fibroblasts are heterogenous populations of ‘accessory’ cells that provide structural support for ‘customer’ cell subsets including the endothelium^27^. Remarkably, approximate 63% endothelial cells with isolated CD31^+^ staining in tumors from *Tgfbr2^fl/fl^*PyMT mice were not associated with GP38^+^ fibroblasts (Fig. 3a-3b), whereas extended layers of GP38^+^ cells enclosed the vasculature in tumors from ThPOK^Cre^*Tgfbr2^fl/fl^*PyMT mice (Fig. 3b). In addition to the supportive cellular compartment, acellular components including extracellular matrix proteins regulate vascular integrity^28^. Associated with the aberrant vessel patterning, the vessel basement membrane proteins collagen IV and fibronectin were fragmented and disoriented in tumors from *Tgfbr2^fl/fl^*PyMT mice (Fig. 3c). In contrast, both collagen IV and fibronectin were highly connected and colocalized with the endothelium in tumors from ThPOK^Cre^*Tgfbr2^fl/fl^*PyMT mice (Fig. 3c). Together, these findings reveal that blockade of TGFβ signaling in CD4^+^ T cells promotes the generation of an organized and mature vasculature in the tumor.

Sprouting angiogenesis is induced in malignant tissues in response to hypoxia and metabolic stresses, which resupplies oxygen and nutrients to a growing tumor^29,30^. Indeed, excessive vessel branching in tumors from *Tgfbr2^fl/fl^*PyMT mice was associated with few hypoxic spots (Fig. 3d). In contrast, approximate 18-fold larger areas were positive for the hypoxic probe in tumors from ThPOK^Cre^*Tgfbr2^fl/fl^*PyMT mice (Fig. 3d). Notably, the hypoxic areas exhibited a circular pattern, and was localized peripheral to the cancer cell death region with the initiating hypoxia and cancer cell death area positioned about 60 μm and 101 μm inward to the adjacent vasculature, respectively (Fig. 3d), which was in agreement with the diffusion range of oxygen in tissue^29,30^. Together, these observations suggest that cancer cell death in ThPOK^Cre^*Tgfbr2^fl/fl^*PyMT mice is caused by severe hypoxia and/or depletion of nutrients, which is enabled by an organized vasculature refractory to the damaging effect of a growing tumor.

### TGFβ represses type 2 immunity to cancer

To define how TGFβRII-deficient CD4^+^ T cells reprogram the cancer microenvironment to fortify vasculature organization, we analyzed tumor-infiltrating T cells from *Tgfbr2^fl/fl^*PyMT and ThPOK^Cre^*Tgfbr2^fl/fl^*PyMT mice. Although the frequencies of tumor-associated CD4^+^Foxp3^+^ Treg cells were not significantly altered, conventional CD4^+^Foxp3^-^ T cells expanded at the expense of CD8^+^ T cells in ThPOK^Cre^*Tgfbr2^fl/fl^*PyMT mice (Extended Data Fig. 6). Importantly, Treg cell-specific ablation of *Tgfbr2* with Foxp3^Cre^ transgenic mice did not affect tumor progression (Fig. 1f), supporting that TGFβ targets T helper (Th) cells to promote cancer immune evasion.

As a pleiotropic regulator of helper T cell responses, TGFβ potently suppresses Th1 and Th2 cell differentiation characterized by expression of IFN-γ and IL-4, respectively^19^. Indeed, increased frequencies of CD4^+^Foxp3^-^ cells from tumor-draining lymph nodes and tumor tissues of ThPOK^Cre^*Tgfbr2^fl/fl^*PyMT mice produced IFN-γ and IL-4 (Fig. 4a and data not shown). In transplantation models of murine cancer, recent studies have shown that Th1 cells and IFN-γ promote pericyte coverage of the endothelium as well as vessel regression^31,32^. To interrogate the function of IFN-γ in the transgenic breast cancer model, we crossed ThPOK^Cre^*Tgfbr2^fl/fl^*PyMT mice onto the IFN-γ-deficient background. Unexpectedly, the repressed tumor growth phenotype was unaffected in the absence of IFN-γ (Fig. 4b). In addition, IFN-γ deficiency did not affect the enhanced effector/memory phenotype of TGFβRII-deficient CD4^+^ T cells in the tumor-draining lymph nodes, or their expansion in the tumor (Extended Data Fig. 7a-7b), while CD8^+^ T cell numbers were reduced (Extended Data Fig. 6 and 7b). Furthermore, extravascular deposition of fibrinogen and the bluntly ended vasculature were observed in PyMT mice on the IFN-γ-deficient background, but were suppressed in *Ifng*^-/-^ThPOK^Cre^*Tgfbr2^fl/fl^*PyMT mice (Extended Data Fig. 7c). Importantly, in the absence of IFN-γ, the severe hypoxia response observed in ThPOK^Cre^*Tgfbr2^fl/fl^*PyMT mice was preserved with clustered CC3-positive cells distributed to the inner circle of the hypoxic region (Extended Data Fig. 7d). These observations exclude IFN-γ as a mediator of cancer environment reprogramming by TGFβRII-deficient CD4^+^ T cells.

To investigate type 2 immune response in cancer regulation, we crossed ThPOK^Cre^*Tgfbr2^fl/fl^*PyMT mice to the IL-4-deficient background. IL-4 deficiency did not affect the enhanced effector/memory differentiation phenotype of TGFβRII-deficient CD4^+^ T cells in the tumor-draining lymph nodes, but their expansion in the tumor was attenuated (Extended Data Fig. 6 and 8a-8b). In contrast to *Ifng*^-/-^ThPOK^Cre^*Tgfbr2^fl/fl^*PyMT mice, *Il4*^-/-^ ThPOK^Cre^*Tgfbr2^fl/fl^*PyMT mice had widespread extravascular deposition of fibrinogen associated with a torturous and irregularly shaped vasculature (Extended Data Fig. 8c). Furthermore, enhanced hypoxia and increased cancer cell death observed in ThPOK^Cre^*Tgfbr2^fl/fl^*PyMT mice were inhibited in the absence of IL-4 (Fig. 3d and Extended Data Fig. 8d), concomitant with accelerated tumor growth (Fig. 4b). In adoptive T cell transfer experiments, conventional CD4^+^ T cells from ThPOK^Cre^*Tgfbr2^fl/fl^*, but not *Tgfbr2^fl/fl^*, *Il4*^-/-^ or *Il4*^-/-^ThPOK^Cre^*Tgfbr2^fl/fl^* mice triggered cancer cell death and halted tumor progression in PyMT recipients (Fig. 4c and Extended Data Fig. 9a), revealing an essential function for IL-4 produced by TGFβRII-deficient CD4^+^ T cells in cancer suppression. Depletion of tumor-associated macrophages that are prominently induced in PyMT mice^33^ did not impair the repressed tumor growth phenotype in ThPOK^Cre^*Tgfbr2^fl/fl^*PyMT mice (Extended Data Fig. 9b-9c), suggesting that macrophages are unlikely effectors of TGFβRII-deficient CD4^+^ T cells. Similar to the PyMT model, blockade of TGFβ signaling in CD4^+^ T cells repressed tumor growth in a transplantation model of cancer, which was blunted by neutralization of IL-4, but not IFN-γ (Extended Data Fig. 9d). These observations suggest a critical role for type 2 immunity in mediating the anti-tumor response triggered by TGFβ deficient CD4^+^ T cells.

### A type 2 gene signature stratifies cancer risk

Aside from being the signature Th2 effector cytokine, IL-4 potently regulates T cell responses. To explore the TGFβ-regulated IL-4-dependent gene expression program in helper T cells, we isolated tumor-infiltrating CD4^+^CD25^-^ T cells from *Tgfbr2^fl/fl^*PyMT, ThPOK^Cre^*Tgfbr2^fl/fl^*PyMT, *Il4*^-/-^ PyMT, and *Il4*^-/-^ThPOK^Cre^*Tgfbr2^fl/fl^*PyMT mice, and performed RNAseq experiments. Compared to wild-type T cells, TGFβRII-deficient T cells upregulated and downregulated 919 and 1507 genes, respectively, among which 238 and 879 were dependent on IL-4 (Supplementary Table 1). We were particularly interested in the 238 transcripts induced in TGFβRII-deficient T cells in an IL-4-depenent manner, as it might define the functional effector program involved in cancer suppression. Among the targets were those encoding stimulatory receptors and adhesion molecules such as *Icam1*, *Il1r2*, *Il3ra*, *Tnfrsf18* and *Tnfrs4*, signaling molecules such as *Jak3*, *Map2k3*, *Map3k10*, *Map3k11*, *Mapkapk2*, *Pik3c2b*, *Sh3bp2* and *Zap70*, calcium, glucose and amino acid transporters including *Orai1*, *Slc2a1*, *Slc2a8* and *Slc7a5* as well as glycolytic enzymes including *Hk2*, *Ldha*, *Pgk1* and *Pkm* (Fig. 5a), which may collectively promote expansion of TGFβRII-deficient CD4^+^ T cells in tumor (Extended Data Fig. 6 and 8b). In addition, a number of transcripts encoding secreted molecules were upregulated in TGFβ deficient T cells in an IL-4-depenent manner, which comprised of Th2 cytokines *Il13*, *Il4* and *Il5* as well as type 2 effector molecules including *Areg*, *Metrnl*, *R3hdml* and *Retnlg* (Fig. 5a), in agreement with potent Th2 cell-inducing activities of IL-4. Indeed, transcripts including *Batf*, *Bhlhe40* and *Pparg* encoding transcription factors resided in major regulatory nodes of T cell activation and Th2 cell differentiation^34^ were as well induced in an IL-4-dependent manner in TGF RII-deficient T cells (Fig. 5a). Collectively, these observations suggest that IL-4 promotes cancer immunity by engaging a positive feedback loop of T cell regulation, although IL-4 targets other than T cells may also be involved.

To determine whether a similar type 2 immune response suppresses tumor development in patients, we refined the IL-4-dependent upregulated gene list by excluding transcripts that did not have human orthologs. We further utilized FPKM filters of non-hematopoietic cancer cell line samples as well as tumor-infiltrating myeloid cells and type 1 lymphocytes, and derived a putative human type 2 immune signature, consisting of 28 genes (Fig. 5b). Notably, when applied to the bulk RNAseq datasets from the TCGA database, the signature stratified patients for survival probability (Fig. 5c). In fact, clinically indolent cancers such as pheochromocytoma and paraganglioma (PCPG) were among the cancer types with highest gene signature value, while aggressive cancers including esophageal carcinoma (ESCA) had low score (Extended Data Fig. 10a). Among breast invasive carcinoma (BRCA), estrogen receptor-, progesterone receptor- or HER2-positive types, collectively grouped as non triple-negative (non-TN), had a higher signature score than TN BRCA (Extended Data Fig. 10a), with its signature value further differentiating patients for survival probability (Extended Data Fig. 10b). Likewise, clinically less aggressive lower grade glioma (LGG) or kidney chromophobe (KICH) had higher signature scores than ontologically related glioblastoma multiforme (GBM) or kidney renal clear cell carcinoma (KIRC), respectively (Extended Data Fig. 10a), and their signature values further stratified patients for better survival probability (Extended Data Fig. 10b). Histological analysis revealed that compared to KIRC, KICH had a more organized vasculature with discernable stromal demarcation and less isolated CD31^+^ endothelial cell staining (Extended Data Fig. 10c). These findings imply a prominent role for type 2 immunity in restraining sprouting angiogenesis and tumor progression in subsets of cancer patients.

## Discussion

As an ancestral cell growth factor family member, TGFβ exerts pleiotropic regulatory functions on numerous pathophysiological processes including carcinogenesis and immune responses^14,15,35,36^. In transplantation models of cancer, TGFβ suppression of tumor immunity has been attributed to the inhibition of tumor-reactive CTLs^20,21,37,38^. Herein, in a sporadic breast cancer model, we provided genetic evidence for Th2 cells, but not CTLs, as a direct target of TGFβ-dependent tumor immune evasion. Using the same tumor model on a different genetic background, Th2 cells were suggested to promote cancer metastasis^39^. How Th2 cells might be involved in triggering different cancer outcomes remains unknown, but could be due to contextual functions of IL-4 manifested with the blockade of TGFβ signaling in Th cells. Nonetheless, the strong anti-tumor phenotype observed in our study challenges the view that type 2 immunity is generally tumor-promoting, but is in agreement with the anti-tumor effects of ectopically administered Th2 cells or overexpression of the Th2-promoting epithelial cell alarmin _TSLP_40-42.

Enhanced activation and differentiation of TGFβRII-deficient CD4^+^ T cells in the tumor-draining lymph nodes and their preferential stromal localization in the tumor tissue are supported by our previous findings that TGFβ1 produced by CD4^+^ T cells, but not cancer cells, suppresses the anti-tumor immune response^18,43,44^; yet, the nature of antigens and accessory signals in promoting Th2 cell differentiation remain to be determined. The exact type 2 effector mechanisms to preserve an ordered vasculature are also open for future investigation. In a rat corneal neovascularization model, administration of IL-4 potently suppresses angiogenesis^45^. Furthermore, “tumor-promoting inflammation”, considered as a cancer enabling characteristic^46^, can be proangiogenic^47-49^; and the broad tumor immune cell exclusion phenotype observed in ThPOK^Cre^*Tgfbr2^fl/fl^*PyMT mice may as well contribute to vasculature organization consequent to the inflammation-resolving activities of Th2 cells. Notably, a similar tumor immune cell exclusion phenotype was observed in a subset of human breast cancer patients^50^, which is associated with dominant CD4^+^ T cell responses and, importantly, predicts better patient survival in line with our type 2 immune signature studies. Thus, the cancer environment-targeted type 2 immune response could represent an evolutionarily conserved cancer defense strategy. Manipulation of such a host-centric cancer immunity pathway may define a novel cancer immunotherapy approach.

## Methods

### Mice

*CD8^-/-^*, *Ifng^-/-^*, *Il4^-/-^* and YFP mice were purchased from the Jackson Laboratory. ThPOK^Cre^ mice were provided by Dr. Ichiro Taniuchi^1^. CD8^Cre^, Foxp3^Cre^, *Tgfrbr2^fl/fl^* and MMTV-PyMT (PyMT) mice were maintained in the laboratory as previously described^2-4^. All mice were backcrossed to the C57BL/6 background, and maintained under specific pathogen-free conditions. Animal experimentation was conducted in accordance with procedures approved by the Institutional Animal Care and Use Committee of Memorial Sloan Kettering Cancer Center.

### Tumor measurement.

Starting from 13-week of age, mammary tumors in female PyMT mice were measured weekly with a caliper. For macrophage or CD4^+^ T cell depletion, 100 μg anti-Csf1r or 500 μg anti-CD4 (BioXcell) were intraperitoneally injected into tumor-bearing mice twice a week. Tumor burden was calculated using the equation [(LxW^2^) x (π/6)], in which L and W denote length and width. Total tumor burden was calculated by summing up individual tumor volumes of each mouse with an end-point defined when total burden reached 3,000 mm^3^ or one tumor reached 2,000 mm^3^, typically around 23-week of age. MC38 cells were a gift from Dr. Jedd Wolchok^5^. 0.5 million MC38 cells were transplanted subcutaneously into *Tgfbr2^fl/fl^* and ThPOK^Cre^*Tgfbr2^fl/fl^* mice followed or not by twice a week treatment with 100 μg anti-IFN-γ (XMG1.2) or anti-IL-4 (11B11). Tumor growth was monitored. Researchers were blinded to genotypes of mice during measurements.

### Leukocyte isolation.

Single-cell suspensions were prepared from lymph nodes by tissue disruption with glass slides. The dissociated cells were passed through 70 μm filters and pelleted. Tumor-infiltrating immune cells were isolated from mammary tumors as previously described6. Briefly, tumor tissues were minced with a razor blade, and digested in 280 U/mL Collagenase Type 3 (Worthington Biochemical) and 4 μg/mL DNase I (Sigma) in HBSS at 37°C for 1 h and 15 min with periodic vortex every 20 min. Digested tissues were passed through 70 μm filters and pelleted. Cells were resuspended in 40% Percoll (Sigma) and layered above 60% Percoll. Sample was centrifuged at 1,900 g at 4°C for 30 min without brake. Cells at interface were collected and used for flow cytometry analysis or sorting.

### Flow cytometry.

Fluorochrome-conjugated or biotinylated antibodies against mouse CD25 (PC61.5), CD4 (RM4-5), CD8 (53-6.7), CD44 (IM7), CD62L (MEL-14), Foxp3 (FJK-16s), IFN-γ (XMG1.2), IL-4 (BVD6-24G2), NK1.1 (PK136), PD-1 (RMP1-130), TCRγδ (eBioGL3) and TCRβ (H57-595) were purchased from eBioscience. Antibody against mouse CD45 (clone 30-F11) was purchased from BD Biosciences. Antibody against granzyme B (GB11) was obtained from Invitrogen. All antibodies were tested with their respective isotype controls. Cell-surface staining was conducted by incubating cells with antibodies for 30 min on ice in the presence of 2.4G2 mAb to block FcγR binding. For Foxp3 and granzyme B staining, a transcription factor-staining kit (Tonbo Biosciences) was used. To assess cytokine production, T cells were stimulated with 50 ng/mL phorbol 12-myristate 13-acetate (Sigma), 1 mM ionomycin (Sigma) in the presence of Golgi-Stop (BD Biosciences) for 4 h at 37°C as previously described^7^. T cells were subsequently stained for cell surface markers before intracellular cytokine staining. All data were acquired using an LSRII flow cytometer (Becton Dickinson) and analyzed with FlowJo software (Tree Star, Inc.).

### Cell sorting, RNA extraction and sequencing.

T cells were FACS sorted to Trizol LS (Invitrogen) and snap frozen in liquid nitrogen. RNA was prepared with a miRNeasy Mini Kit according to the manufacturer’s instructions (Qiagen), and subject to quality control by Agilent BioAnalyzer. 0.7-1 ng total RNA with an integrity index from 8.3 to 9.9 was amplified using the SMART-Seq v4 Ultra Low Input RNA Kit (Clontech), with 12 cycles of amplification. 2.7 ng of amplified cDNA was used to prepare libraries with the KAPA Hyper Prep Kit (Kapa Biosystems KK 8504). Samples were barcoded and used for 50bp/50bp paired end runs with the TruSeq SBS Kit v4 (Illumina) on a HiSeq 2500 sequencer. An average of 45 million paired reads were generated per sample. The percentage of mRNA bases per sample ranged from 75% to 81%.

### Transcriptome analysis of differentially expressed genes.

The raw sequencing FASTQ files were aligned against the mm10 assembly by STAR8. Gene level count values were computed by the summarizeOverlaps function from the R package “GenomicAlignments” with mm10 KnownGene as the base gene model for mouse samples9,10. The Union counting mode was used and only mapped paired-reads were considered. FPKM (Fragments Per Kilobase Million) values were then computed from gene level counts by using fpkm function from the R package DESeq211. Differentially expressed gene analysis was performed through the R DESeq2 package. Given the raw count data and gene model used, DESeq2 normalized the expression raw count data by sample specific size factor and took specified covariates into account while testing for genes found with significantly different expression between 2 sample groups. The comparisons included ThPOKCreTgfbr2fl/flPyMT (KO) group vs. Tgfbr2fl/flPyMT (WT) group, KO group vs. Il4-/-ThPOKCreTgfbr2fl/flPyMT (Il4-/-KO) group, Il4-/-KO group vs. Il4-/-Tgfbr2fl/flPyMT (Il4-/-) group, and Il4-/-KO group vs. WT group. The “log2FoldChange” column is the log2 value of the ratio between target sample group and the reference sample group. Differentially expressed genes were included if passed the following filters: baseMean > 50; Log2Foldchange > 0.9 or < −0.9 (depending on the desired direction) and P value < 0.11. For heatmap generation, the mean expression FPKM value in each mouse sample group was calculated. The ratio between the sample group specific mean expression FPKM value and the average of 4 sample group mean expression FPKM values was used and plotted in the heatmap through the R heatmap package.

### Gene signature building.

The differentially expressed genes in the same regulation direction from comparing ThPOK^Cre^*Tgfbr2^fl/fl^*PyMT (KO) group vs. *Tgfbr2^fl/fl^*PyMT (WT) group and KO group vs. *Il4^-/-^*ThPOK^Cre^*Tgfbr2^fl/fl^*PyMT (*Il4^-/-^*KO) group were subjected for human homologous gene search. The corresponding human homologous gene of mouse gene was identified through referencing the Mouse Genome Informatics (MGI, http://www.informatics.jax.org/) database as well as the NCBI Orthologs database (ftp://ftp.ncbi.nlm.nih.gov/gene/DATA/gene_orthologs.gz). Any mouse gene with missing human homolog was excluded from building the human gene signature. Any human gene found with expression greater than 25 FPKM in any non-hematopoietic Cancer Cell Line sample from Broad Institute (https://portals.broadinstitute.org/ccle) or with expression greater than 25 FPKM in any follow cytometry-sorted human RCC-infiltrated myeloid and type 1 lymphocyte samples were excluded from the signature building.

### Gene signature applied to TCGA Pan cancer cohort.

Single-Sample Gene Set Enrichment Analysis (ssGSEA) was done by applying the gene signature against the FPKM expression values of TCGA Pan cancer samples through the R package “GSVA” which estimates the signature enrichment score for each of RNASeq sample12. The TCGA Pan cancer cohort data were downloaded from the GDC (https://gdc.cancer.gov/aboutdata/publications/pancanatlas) including RNA-seq expression matrix as well as the clinical data resource file.

### Survival analysis K-M plot.

Survival analyses and curves were performed and generated according to the Kaplan–Meier method using SAS v.9.4 (SAS Institute Inc., Cary, North Carolina). The log-rank test was used to determine the statistical significance of the overall survival (OS) distributions between signature high and signature low groups defined by the median signature ssGSEA score in TCGA Pan cancer cohort.

### Immunofluorescence staining.

Antibodies against mouse CD31 (MEC13.3), CD3 (17A2) and GP38 (8.1.1) were purchased from Biolegend. Antibody against mouse CD45 (30-F11) was from BD Pharmingen. Antibodies against mouse Col IV (Cat. #2150-1470), Fibrinogen (Cat. #4440-8004) and NG2 (Cat. #AB5320) were obtained from Bio-rad. Antibody against mouse Fibronectin (Cat. #AB2033) was from EMD. Antibody against mouse Cleaved caspase 3 (Cat. #9661S) was purchased from CST. Antibodies against mouse E-Cadherin (DECMA-1) and Ki67 (SolA15) were obtained from eBioscience. Antibodies against human E-Cadherin (Cat ab40772) and CD31 (Cat #ab76533) were purchased from Abcam. Hypoxia detection kit was purchased from Hypoxyprobe. Mouse tumor tissues with comparable size from control and experimental groups were frozen in O.C.T. medium (Sakura Finetek USA) and sectioned at the thickness of 10 μm, before they were fixed and stained with antibodies or probes. Human clear cell and chromophobe renal cell carcinoma paraffin sections with 5 μm thickness were dewaxed, rehydrated, blocked and stained with antibodies. Slide sections were mounted with VECTORSHIELD anti-fade mounting media (Vector Laboratories) and scanned by Pannoramic Digital Slide Scanners (3DHISTECH LTD). Immunofluorescence images were analyzed with CaseViewer and Fiji software, and further processed with Adobe Photoshop and Illustrator software.

### T cell activation and adoptive T cell transfer.

CD4+CD25-T cells isolated from *Tgfbr2fl/fl, ThPOKCreTgfbr2fl/fl, Il4-/-Tgfbr2fl/fl and Il4-/-ThPOKCreTgfbr2fl/fl* mice were activated with plate-bound anti-CD3 (5 μg/mL) and soluble anti-CD28 (1 μg/mL) for 24 h. One million T cells were intravenously transferred to tumor-bearing PyMT mice on a weekly basis.

### Sulfo-NHS-biotin labeling.

Deeply anesthetized mice were perfused with 10 mL Sulfo-NHS-LC-Biotin (Thermo Scientific; 21335) dissolved in PBS at 0.3 mg/mL, followed by 10 mL 2% PFA in PBS^13^.

### Statistical analysis

Related to Fig. 5 and Supplementary Table 1, differentially expressed genes were compared between tumor-infiltrating CD4^+^CD25^-^ T cells from *Tgfbr2^fl/fl^*PyMT, ThPOK^Cre^*Tgfbr2^fl/fl^*PyMT, *Il4^-/-^Tgfbr2^fl/fl^*PyMT and *Il4^-/-^*ThPOK^Cre^*Tgfbr2^fl/fl^*PyMT mice. A gene list was generated if passing the filters mentioned above. All statistical measurements are displayed as mean ± SEM. For comparisons, unpaired student t test, two-tailed was conducted using GraphPad Prism software; for paired distance comparisons, paired t-test was conducted using GraphPad Prism software. For tumor growth, 2-way ANOVA was performed using GraphPad Prism software.

## Acknowledgements

We thank members of the M.O.L. laboratory for helpful discussions. This work was supported by a Howard Hughes Medical Institute Faculty Scholar Award (M.O.L.), an award from Mr. William H. and Mrs. Alice Goodwin and the Commonwealth Foundation for Cancer Research and the Center for Experimental Therapeutics at Memorial Sloan Kettering Cancer Center (M.O.L.) and a Cancer Center Support Grant (P30 CA08748). B.G.N. and M.H.D. are recipients of F31 CA210332 and F30 AI29273-03 awards from National Institutes of Health. C.C. and S.L. are Cancer Research Institute Irvington Fellows supported by the Cancer Research Institute. E.G.S. is a recipient of a Fellowship from the Alan and Sandra Gerry Metastasis and Tumor Ecosystems Center of MSKCC. MSKCC has filed a patent application with the U.S. Patent and Trademark Office directed toward methods and compositions of targeting CD4^+^ T helper cell TGFβ signaling for caner immunotherapy.

## Author Contributions

M.L. and M.O.L. were involved in all aspects of this study, including planning and performing experiments, analysis and interpretation of data and writing the manuscript. F.K. and T.A.C. processed and analyzed all sequencing data and wrote the manuscript. I.T. provided the key mouse line. Y.C., J.J.H. and A.A.H. provided human RCC specimen. K.J.C., D.K., B.G.N., W.S., C.C., M.H.D., E.G.S. and S.L. assisted with mouse colony management and performed experiments.

## Extended Data Figure Legends

**Extended Data Figure 1.**
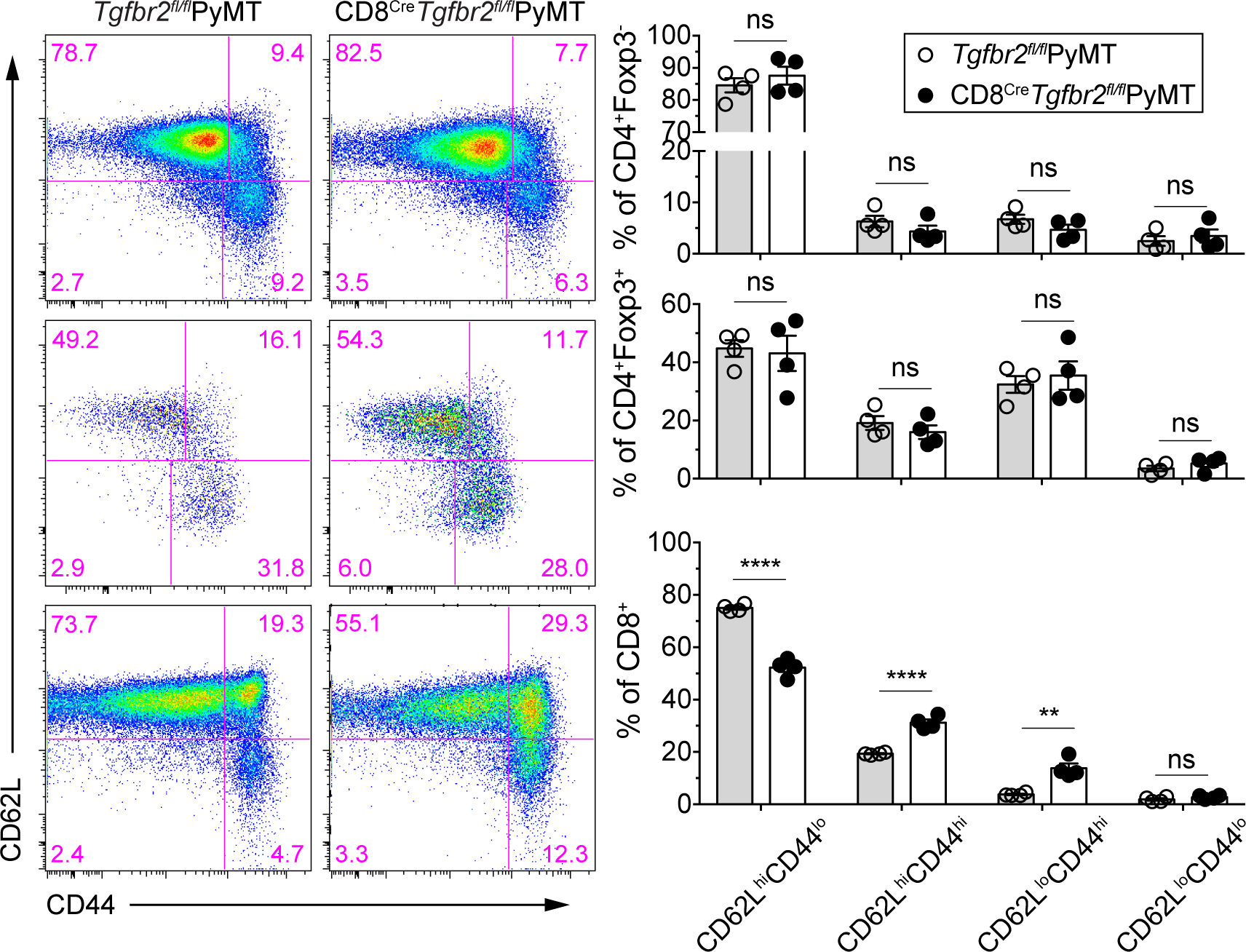
**Blockade of TGF**β **signaling in CD8^+^ T cells promotes effector/memory differentiation** Representative flow cytometry plots of CD62L and CD44 expression and statistical analyses of the gated populations among CD4^+^Foxp3^-^ T cells (top panel), CD4^+^Foxp3^+^ T cells (middle panel) and CD8^+^ T cells (bottom panel) from the tumor-draining lymph nodes of *Tgfbr2^fl/fl^*PyMT and CD8^Cre^*Tgfbr2^fl/fl^*PyMT mice. All statistical data are shown as mean ± SEM. **: P<0.01; ****: P<0.0001; and ns: not significant.

**Extended Data Figure 2.**
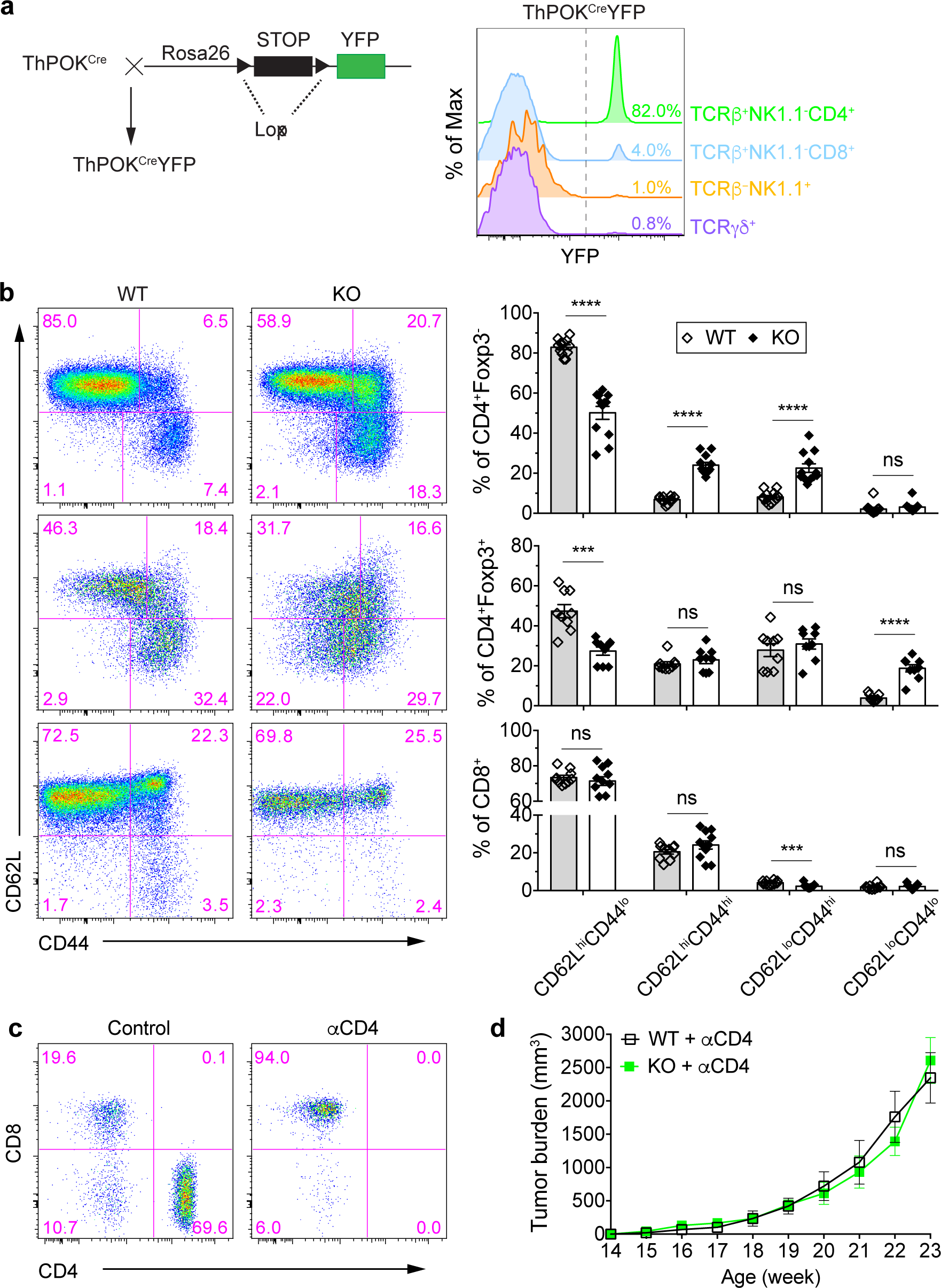
Blockade of TGFβ signaling in CD4^+^ T cells promotes effector/memory differentiation and accounts for reduced tumor growth in ThPOK^Cre^*Tgfbr2^fl/fl^*PyMT mice **a,** A breeding scheme of ThPOK^Cre^ mice with YFP mice and representative flow cytometry plots of YFP expression in lymph node TCRβ^+^NK1.1^-^CD4^+^, TCRβ^+^NK1.1^-^CD8^+^, TCRβ^-^NK1.1^+^ and TCRγδ^+^ T cells isolated from ThPOK^Cre^YFP mice. **b,** Representative flow cytometry plots of CD62L and CD44 expression and statistical analyses of the gated populations among CD4^+^Foxp3^-^ T cells (top panel), CD4^+^Foxp3^+^ T cells (middle panel) and CD8^+^ T cells (bottom panel) from the tumor-draining lymph nodes of *Tgfbr2^fl/fl^*PyMT (wild-type, WT) and ThPOK^Cre^*Tgfbr2^fl/fl^*PyMT (knockout, KO) mice. **c**, Representative flow cytometry plots of CD4 and CD8 expression in T cells isolated from control mice or mice treated with anti-CD4 (αCD4). **d**, Tumor measurements of WT (n=3) and KO (n=3) mice treated with αCD4. All data are shown as mean ± SEM. ***: P<0.001; ****: P<0.0001; and ns: not significant.

**Extended Data Figure 3.**
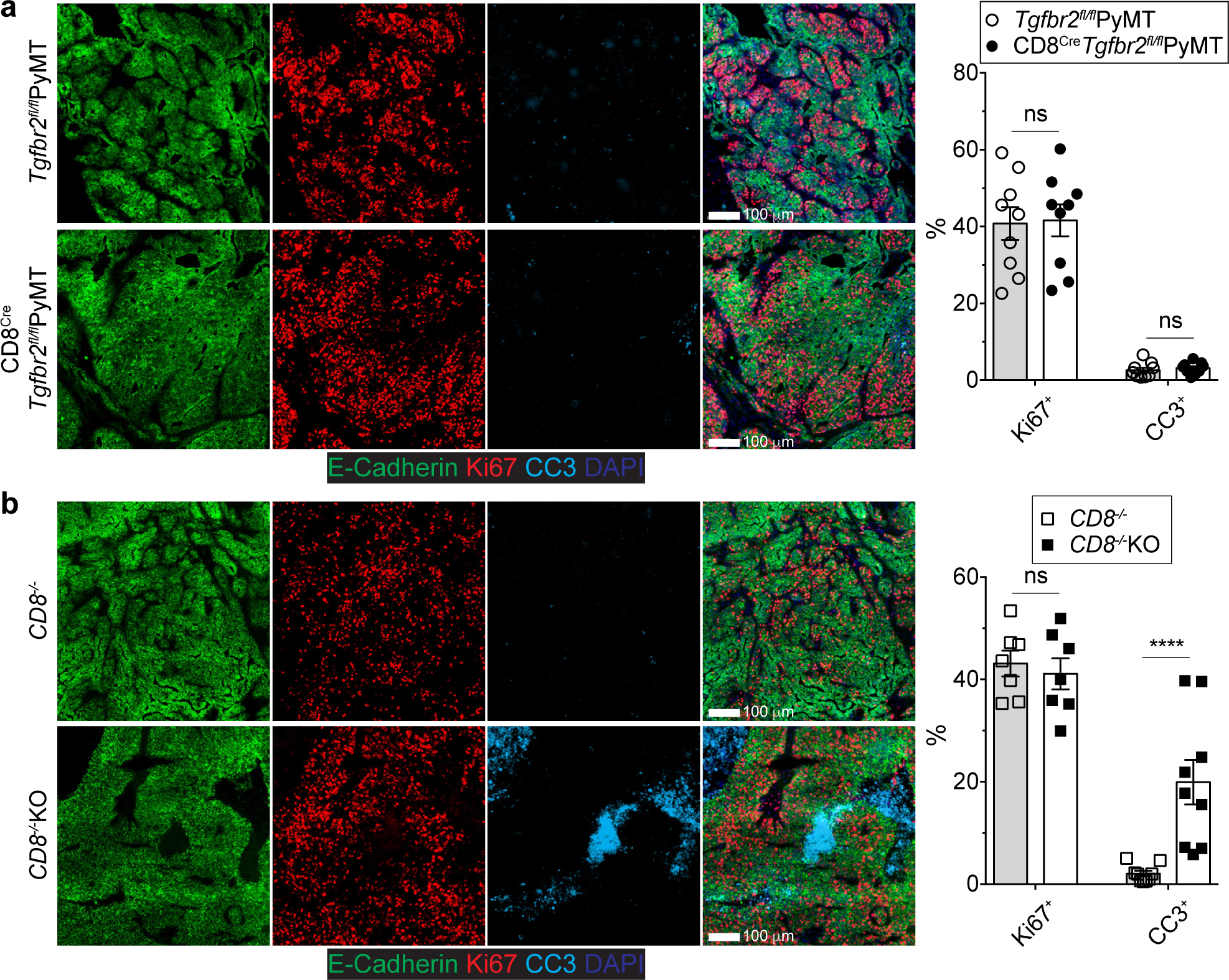
CD8 deficiency does not affect cancer cell death triggered by **blockade of TGF**β **signaling in CD4^+^ T cells a,** Representative immunofluorescence images and statistical analyses of E-Cadherin (green), Ki67 (red) and cleaved Caspase 3 (CC3, cyan) e xp r e s s i o n in mammary tumor tissues from 23-week-old *Tgfbr2^fl/fl^*PyMT and CD8^Cre^*Tgfbr2^fl/fl^*PyMT mice. The percentage of Ki67^+^E-Cadherin^+^ cells over total E-Cadherin^+^ epithelial cells was calculated from multiple 0.02 mm^2^ regions (n=9). The percentage of CC3^+^ areas over total E-Cadherin^+^ areas was calculated from multiple 0.02 mm^2^ regions (n=9). **b,** Representative immunofluorescence images and statistical analyses of E-Cadherin (green), Ki67 (red) and cleaved Caspase 3 (CC3, cyan) e xp r e s s i o n in mammary tumor tissues from 23-week-old *CD8^-/-^Tgfbr2^fl/fl^*PyMT (*CD8^-/-^*) and *CD8^-/-^* ThPOK^Cre^*Tgfbr2^fl/fl^*PyMT (*CD8^-/-^* knockout, *CD8^-/-^*KO) mice. The percentage of Ki67^+^E-Cadherin^+^ cells over total E-Cadherin^+^ epithelial cells was calculated from multiple 0.02 mm^2^ regions (n=9). The percentage of CC3^+^ areas over total E-Cadherin^+^ areas was calculated from multiple 0.02 mm^2^ regions (n=9). All data are shown as mean ± SEM. ****: P<0.0001; and ns: not significant.

**Extended Data Figure 4.**
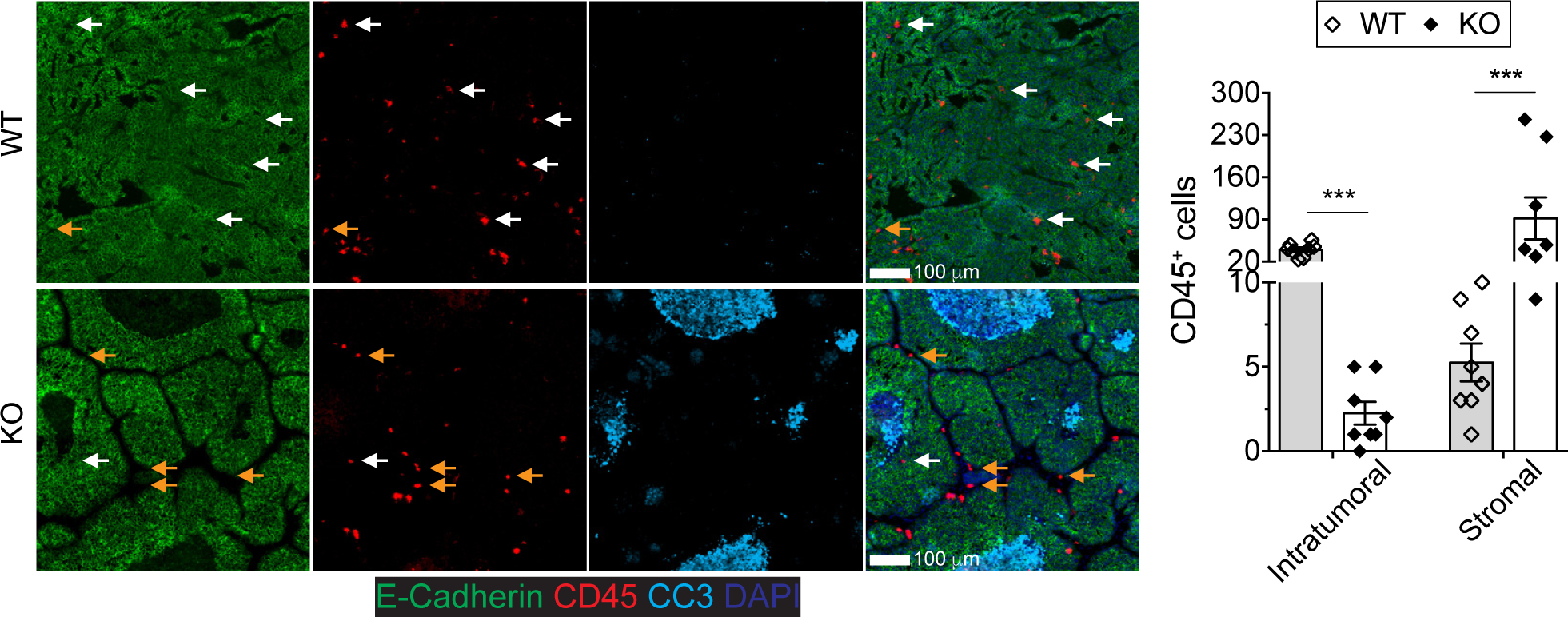
Blockade of TGFβ signaling in CD4^+^ T cells causes leukocyte exclusion from the tumor parenchyma Representative immunofluorescence images and statistical analyses of E-Cadherin (green), CD45 (red) and cleaved Caspase 3 (CC3, cyan) expression in mammary tumor tissues from 23-week-old *Tgfbr2^fl/fl^*PyMT (wild-type, WT) and ThPOK^Cre^*Tgfbr2^fl/fl^*PyMT (knockout, KO) mice. Intratumoral (white arrows) and stromal (yellow arrows) CD45^+^ leukocytes were counted from multiple 0.1 mm^2^ regions (n=8 for WT and KO tumor tissues). All data are shown as mean ± SEM. ***: P<0.001.

**Extended Data Figure 5.**
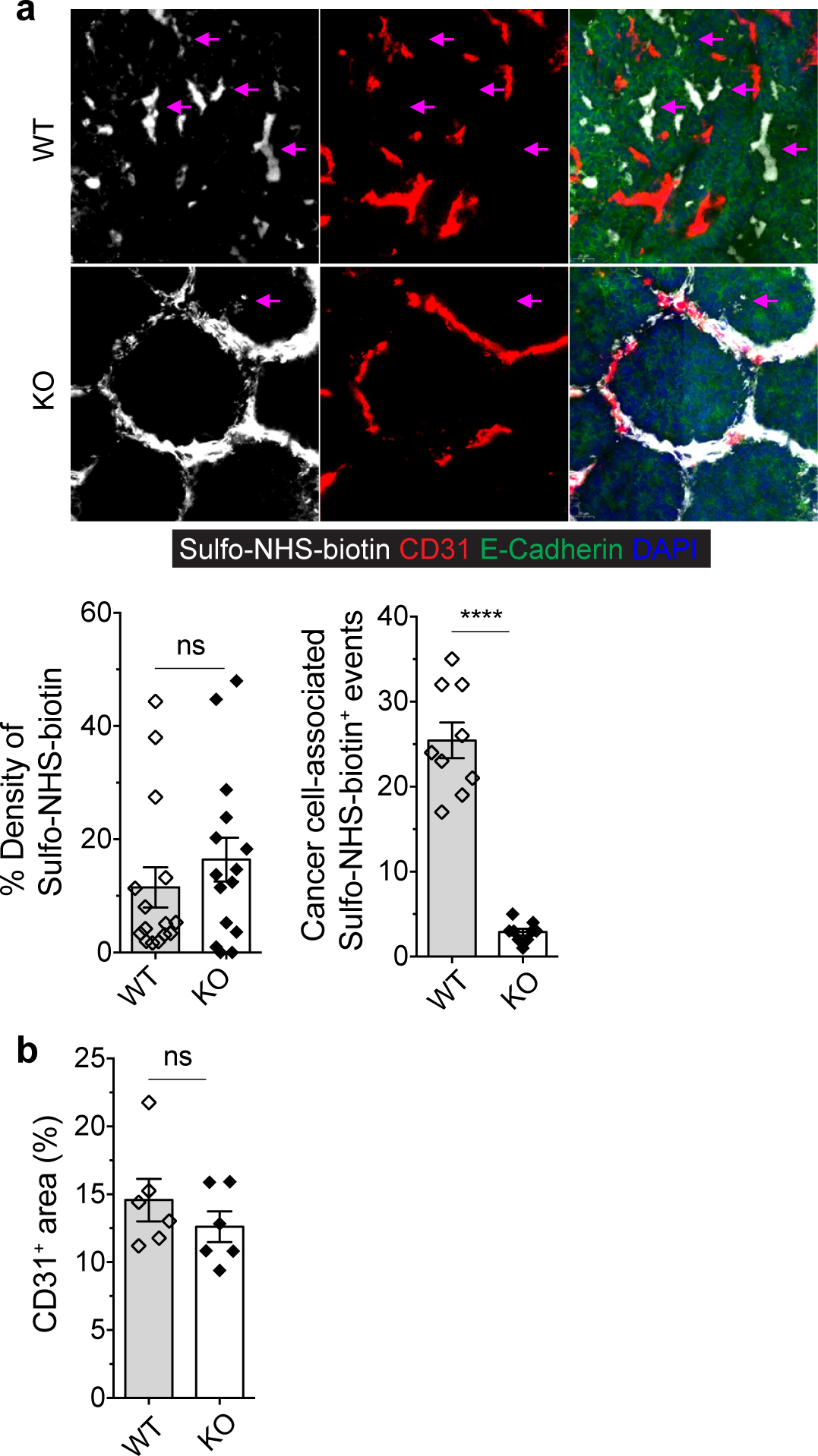
Blockade of TGFβ signaling in CD4^+^ T cells inhibits vasculature leakage, but does not affect vessel density **a**, Representative images of Sulfo-NHS-biotin (white), CD31 (red) and E-Cadherin (green) staining and quantification of Sulfo-NHS-biotin density and cancer cell-associated Sulfo-NHS-biotin^+^ events (magenta arrows) in mammary tumor tissues from 23-week-old *Tgfbr2^fl/fl^*PyMT (wild-type, WT) and ThPOK^Cre^*Tgfbr2^fl/fl^*PyMT (knockout, KO) mice. The percentage of Sulfo-NHS-biotin^+^ areas (n=15) and cancer-cell associated deposition events (n=9) were calculated from multiple 0.8 mm^2^ regions. **b**, Quantification of CD31^+^ areas in multiple 1 mm^2^ regions of mammary tumor tissues from 23-week-old WT and KO mice (n=6). All data are shown as mean ± SEM. ****: P<0.001; and ns: not significant.

**Extended Data Figure 6.**
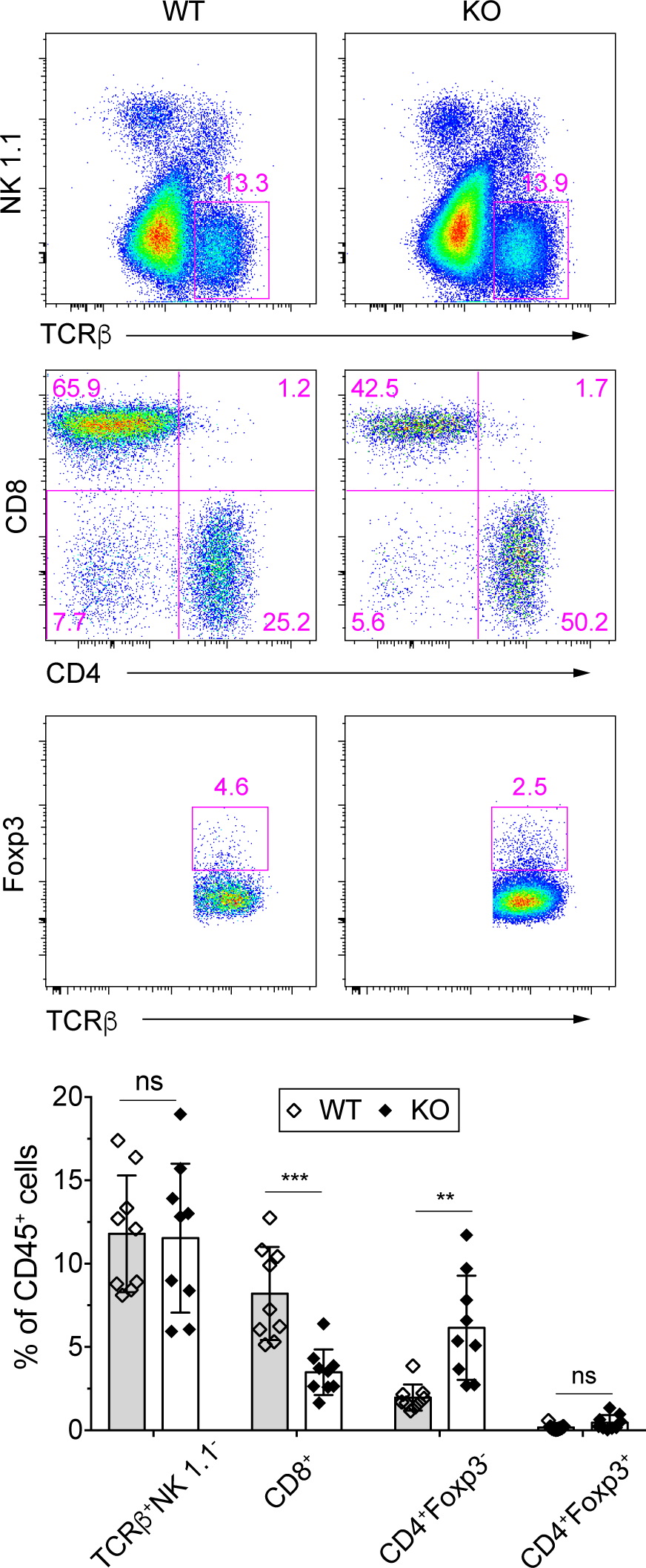
Blockade of TGFβ signaling in CD4^+^ T cells promotes expansion of tumor-infiltrating CD4^+^Foxp3^-^ T cells Representative flow cytometry plots of TCRβ, NK1.1, CD4, CD8 and Foxp3 expression and statistical analyses of the gated populations in tumor-infiltrating leukocytes from 23-week-old *Tgfbr2^fl/fl^*PyMT (WT) and ThPOK^Cre^*Tgfbr2^fl/fl^*PyMT (KO) mice. All data are shown as mean ± SEM. **: P<0.01; ***: P<0.001; and ns: not significant.

**Extended Data Figure 7.**
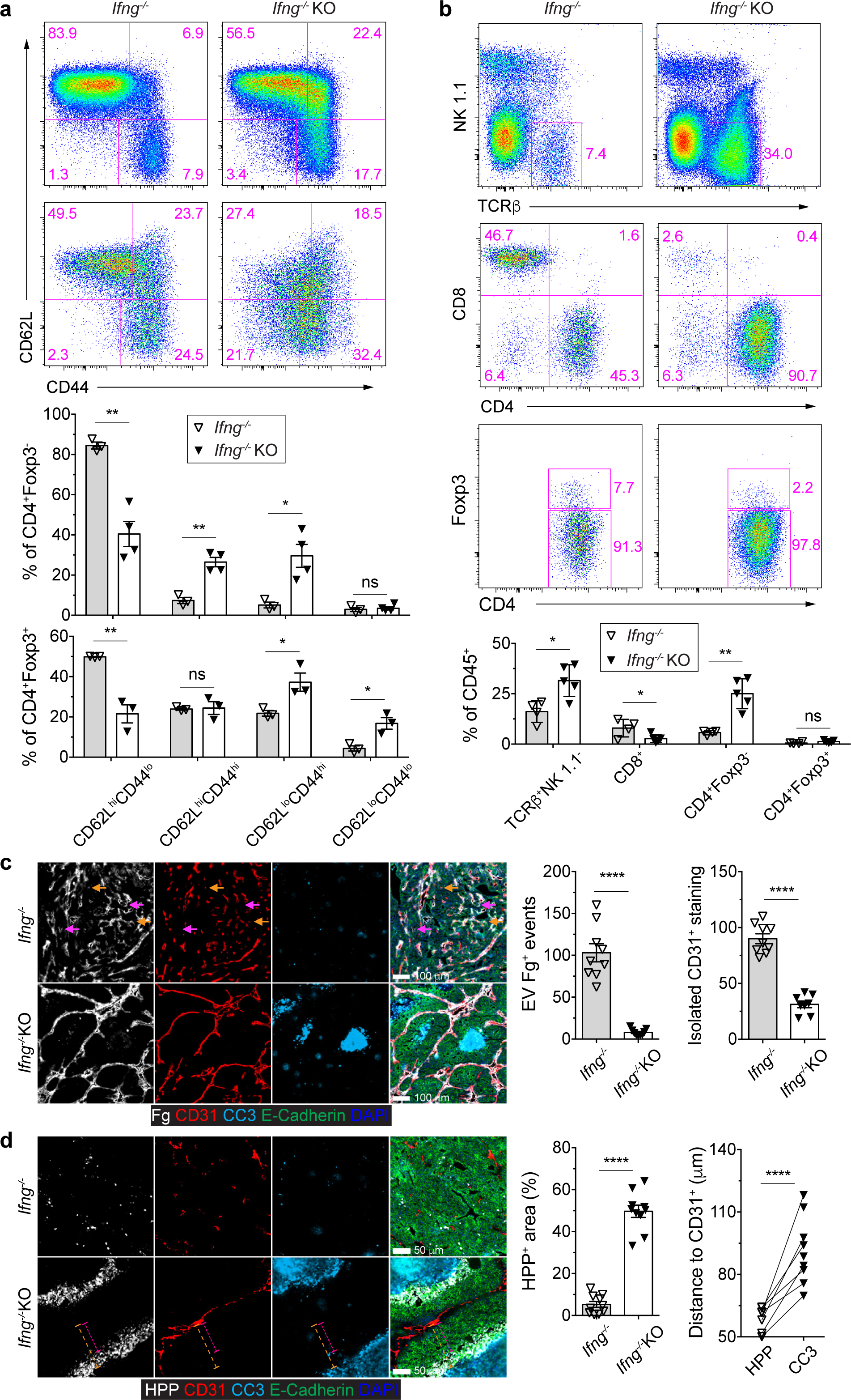
IFN-**γ** deficiency does not impair cancer immunity triggered by blockade of TGFβ signaling in CD4^+^ T cells **a,** Representative flow cytometry plots of CD62L and CD44 expression and statistical analyses of the gated populations among CD4^+^Foxp3^-^ T cells (top panel) and CD4^+^Foxp3^+^ T cells (bottom panel) from *Ifng*^-/-^*Tgfbr2^fl/fl^*PyMT (*Ifng*^-/-^) and *Ifng*^-/-^ThPOK^Cre^*Tgfbr2^fl/fl^*PyMT (*Ifng*^-/-^ knockout, *Ifng*^-/-^KO) mice. **b,** Representative flow cytometry plots of TCRβ, NK1.1, CD4, CD8 and Foxp3 expression and statistical analyses of the gated populations in tumor-infiltrating leukocytes from 23-week-old *Ifng*^-/-^ and *Ifng*^-/-^KO mice. **c,** Representative immunofluorescence images of fibrinogen (Fg, white), CD31 (red), cleaved Caspase 3 (CC3, cyan) and E-Cadherin (green) in comparable individual tumors with sizes around 8×8 mm in length and width from 23-week-old *Ifng*^-/-^ and *Ifng*^-/-^KO mice. Extravascular (EV) Fg deposition events (magenta arrows) were calculated from multiple 1 mm^2^ regions (n=9 for *Ifng*^-/-^ and *Ifng*^-/-^KO tumor tissues). Isolated CD31^+^ staining (yellow arrows) was counted from multiple 1 mm^2^ regions (n=9 for *Ifng*^-/-^ and *Ifng*^-/-^KO tumor tissues). **d,** Representative immunofluorescence images of Hypoxic probe (HPP, white), CD31 (red), cleaved Caspase 3 (CC3, cyan) and E-Cadherin (green) in comparable individual tumors with sizes around 8×8 mm in length and width from *Ifng*^-/-^ and *Ifng*^-/-^KO mice. The percentage of HPP^+^E-Cadherin^+^ regions over E-Cadherin^+^ epithelial regions was calculated from multiple 1 mm^2^ regions (n=9 for *Ifng*^-/-^ and *Ifng*^-/-^KO tumor tissues). The shortest distance of HPP^+^ regions (magenta dashed lines) or CC3^+^ regions (yellow dashed lines) to CD31^+^ endothelial cells was measured in tumor tissues from *Ifng^-/-^*KO mice (n=9). All data are shown as mean ± SEM. *: P<0.05; **: P<0.01; ****: P<0.0001; and ns: not significant.

**Extended Data Figure 8.**
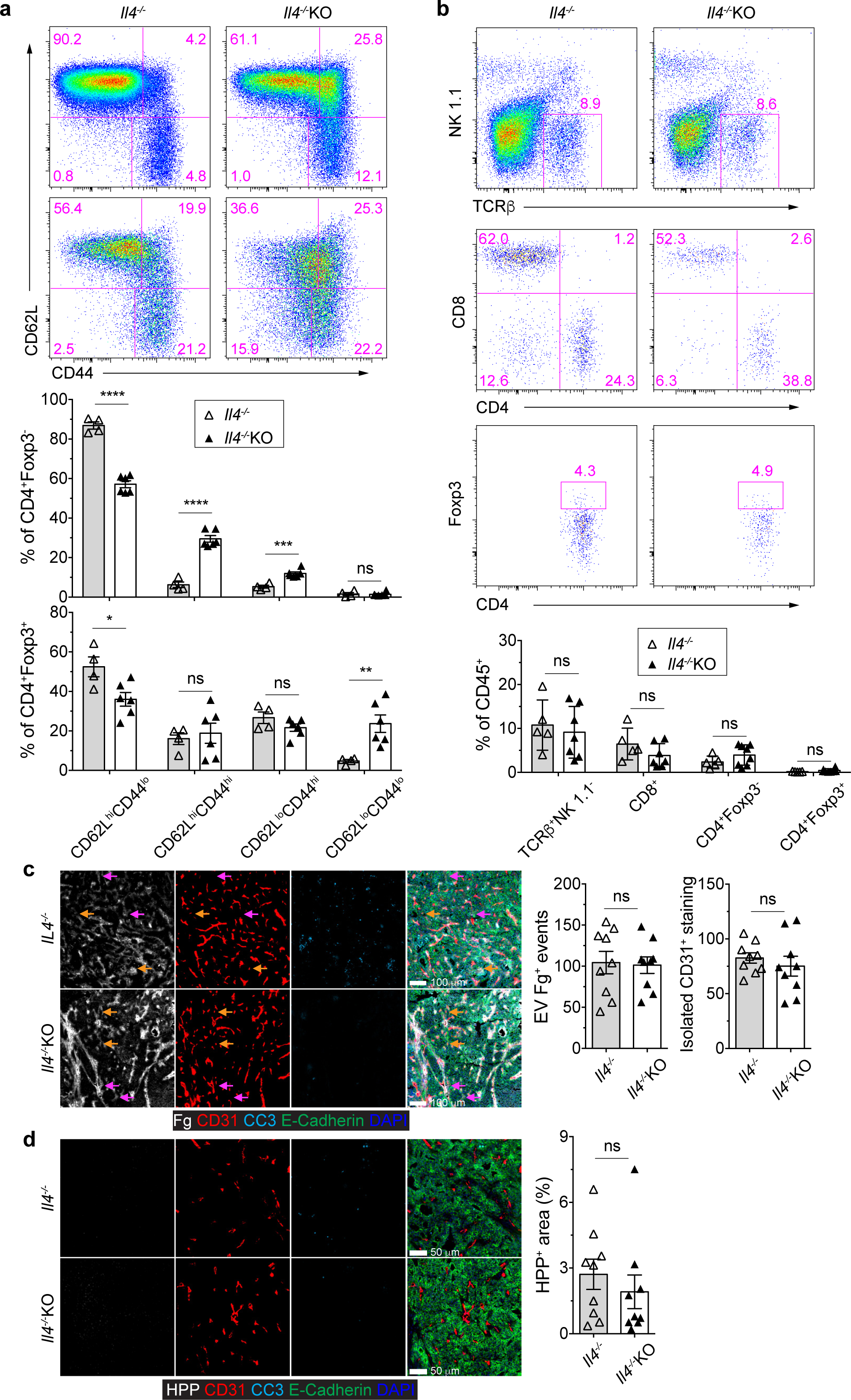
IL-4 deficiency impairs cancer immunity triggered by blockade of TGFβ signaling in CD4^+^ T cells **a,** Representative flow cytometry plots of CD62L and CD44 expression and statistical analyses of the gated populations among CD4^+^Foxp3^-^ T cells (top panel) and CD4^+^Foxp3^+^ T cells (bottom panel) from *Il4*^-/-^*Tgfbr2^fl/fl^*PyMT (*Il4*^-/-^) and *Il4*^-/-^ThPOK^Cre^*Tgfbr2^fl/fl^*PyMT (*Il4*^-/-^ knockout, *Il4*^-/-^KO) mice. **b,** Representative flow cytometry plots of TCRβ, NK1.1, CD4, CD8 and Foxp3 expression and statistical analyses of the gated populations in tumor-infiltrating leukocytes from 23-week-old *Il4*^-/-^ and *Il4*^-/-^KO mice. **c,** Representative immunofluorescence images of fibrinogen (Fg, white), CD31 (red), cleaved Caspase 3 (CC3, cyan) and E-Cadherin (green) in comparable individual tumors with sizes around 8×8 mm in length and width from 23-week-old *Il4^-/-^* and *Il4^-/-^*KO mice. Extravascular (EV) Fg deposition events (magenta arrows) were calculated from multiple 1 mm^2^ regions (n=9 for *Il4^-/-^* and *Il4^-/-^*KO tumor tissues). Isolated CD31^+^ staining (yellow arrows) was counted from multiple 1 mm^2^ regions (n=9 for *Il4^-/-^* and *Il4^-/-^*KO tumor tissues). **d,** Representative immunofluorescence images of Hypoxic probe (HPP, white), CD31 (red), cleaved Caspase 3 (CC3, cyan) and E-Cadherin (green) in comparable individual tumors with sizes around 8×8 mm in length and width from *Il4^-/-^* and *Il4^-/-^*KO mice. The percentage of HPP^+^E-Cadherin^+^ regions over E-Cadherin^+^ epithelial regions was calculated from multiple 1 mm^2^ regions (n=9 for *Il4*^-/-^ and *Il4^-/-^*KO tumor tissues). All data are shown as mean ± SEM. *: P<0.05; **: P<0.01; ***: P<0.001; ****: P<0.0001; and ns: not significant.

**Extended Data Figure 9.**
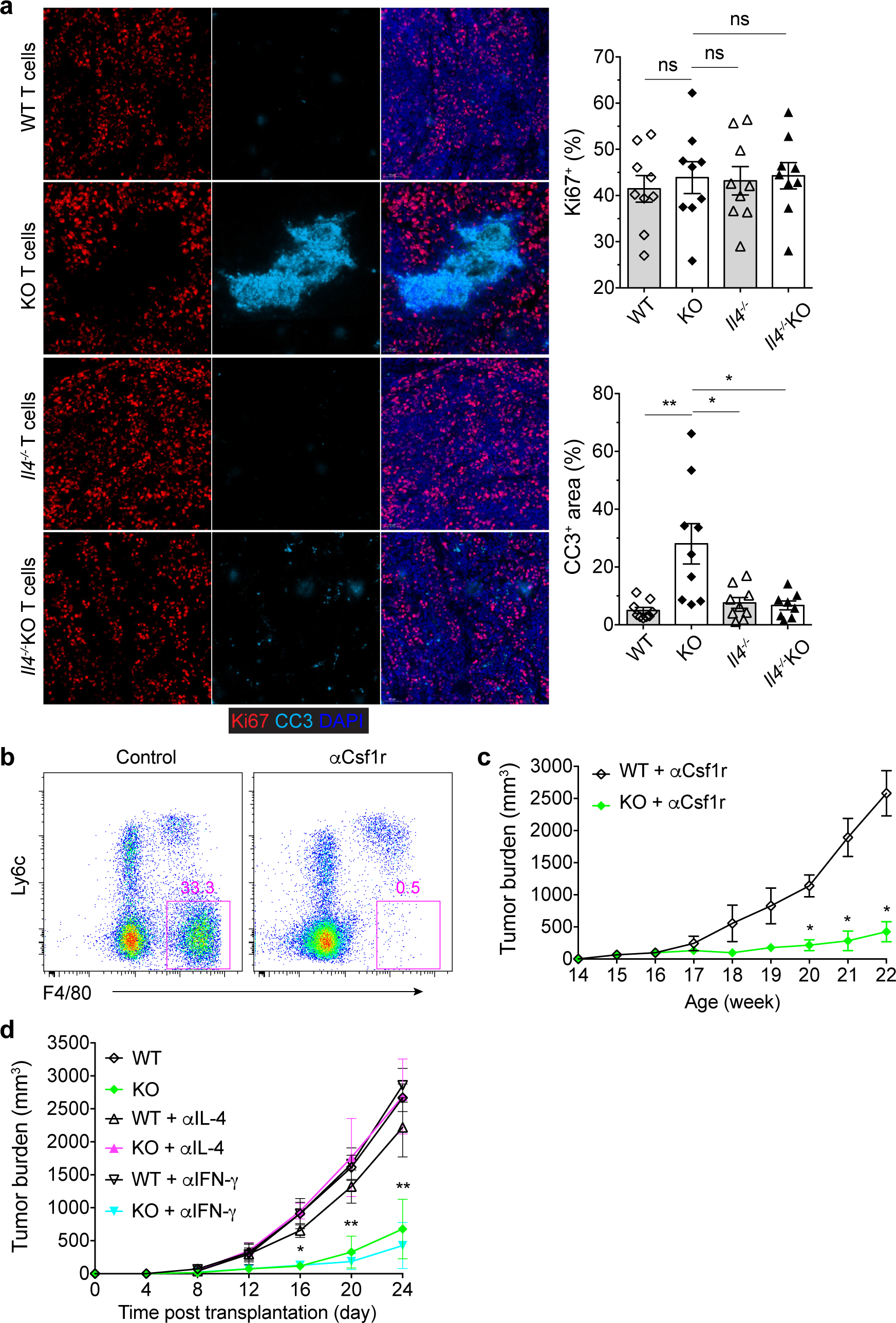
Anti-tumor immunity triggered by TGFβ signaling blockade in CD4^+^ T cells is dependent on IL-4, but not tumor-associated macrophages **a,** 16-week-old PyMT mice bearing 5×5 mm tumors were transferred with the activated CD4^+^CD25^-^ T cells from *Tgfbr2^fl/fl^* (WT), ThPOK^Cre^*Tgfbr2^fl/fl^* (KO), *Il4^-/-^Tgfbr2^fl/fl^* (*Il4^-/-^*) and *Il4^-/-^* ThPOK^Cre^*Tgfbr2^fl/fl^* (*Il4^-/-^*KO) mice on a weekly basis for 6 weeks. Representative immunofluorescence images and statistical analyses of Ki67 (red) and cleaved Caspase 3 (CC3, cyan) expression. The percentage of Ki67^+^E-Cadherin^+^ cells over total E-Cadherin^+^ epithelial cells was calculated from multiple 0.02 mm^2^ regions (n=9). The percentage of CC3^+^ areas over total E-Cadherin^+^ areas was calculated from multiple 0.02 mm^2^ regions (n=9). **b,** Representative flow cytometry plots of F4/80^+^Ly6c^-^ tumor-associated macrophage populations from control mice or mice treated with anti-Csf1r (αCsf1r). **c**, Tumor measurements of WT PyMT (n=3) and KO PyMT (n=3) mice treated with αCsf1r. **d**, Tumor measurements of WT (n=5), KO (n=4), αIL-4-treated WT (n=5), αIL-4-treated KO (n=4), αIFN-γ-treated WT (n=3), and αIFN-γ-treated KO mice (n=3) inoculated with MC38 tumor cells. All data are shown as mean ± SEM. *: P<0.05; **: P<0.01; and ns: not significant.

**Extended Data Figure 10:**
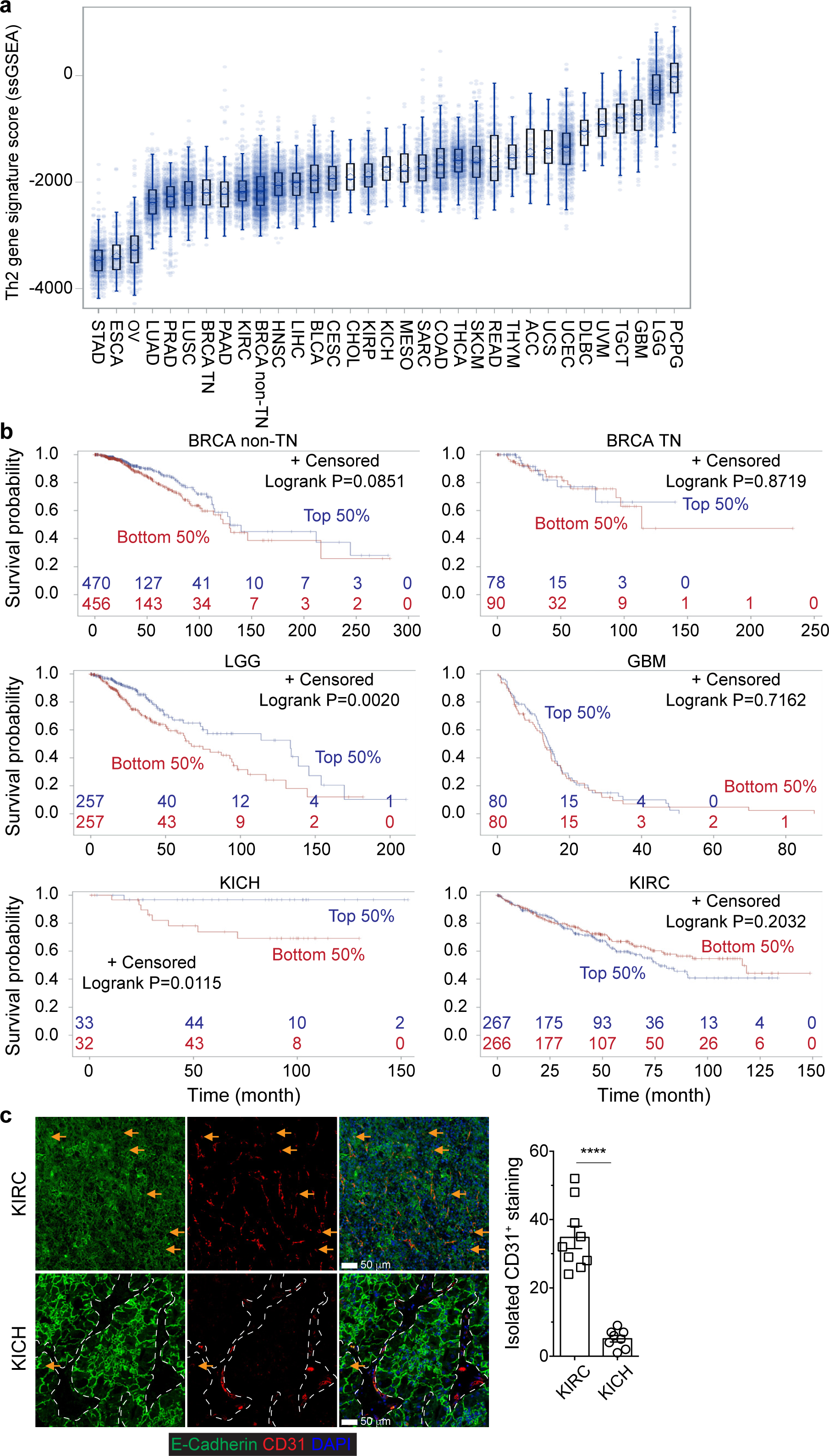
Distribution of a curated Th2 gene expression signature across TCGA Pan cancer patients and its association with the survival probability of selected patient groups and their vasculature status **a,** The human Th2 gene signature enrichment score was estimated and plotted for each RNASeq sample of TCGA Pan cancer patients. **b,** A Th2 gene signature was used to perform survival analysis in TCGA non-triple negative (non-TN) and triple negative (TN) breast invasive carcinoma (BRCA), brain lower grade glioma (LGG) and glioblastoma multiforme (GBM), kidney chromophobe (KICH) and kidney renal clear cell carcinoma (KIRC) patients. The survival curves were plotted for the signature high group (top 50%) and low group (bottom 50%). The corresponding censored patient numbers are included in major time points. **c,** Representative immunofluorescence images and statistical analyses of E-Cadherin (green) and CD31 (red) expression in KICH (4 patients) and KIRC (8 patients) tumor tissues. Isolated CD31^+^ staining was counted from multiple 0.2 mm^2^ regions (n=9). The stromal regions are marked by dotted lines in KICH samples. All data are shown as mean ± SEM. ****: P<0.0001; and ns: not significant.

**Supplementary Table 1**: Differentially expressed genes in tumor-infiltrating CD4^+^CD25^-^ T cells from *Tgfbr2^fl/fl^*PyMT (wild-type, WT), ThPOK^Cre^*Tgfbr2^fl/fl^*PyMT (knockout, KO), *Il4^-/-^ Tgfbr2^fl/fl^*PyMT (*Il4^-/-^*) and *Il4^-/-^*ThPOK^Cre^*Tgfbr2^fl/fl^*PyMT (*Il4^-/-^*KO) mice.

